# Embodied behavioural complexity in a ciliated microorganism

**DOI:** 10.1101/2025.08.20.671109

**Authors:** Alexander K. Boggon, Alasdair D. Hastewell, Jörn Dunkel, Kirsty Y. Wan

**Affiliations:** Living Systems Institute & Department of Mathematics and Statistics, University of Exeter, Stocker Road, Exeter, EX4 4QD, UK; NSF-Simons National Institute for Theory and Mathematics in Biology, Chicago IL, 60611, USA; Department of Mathematics, Massachusetts Institute of Technology, Massachusetts Avenue, Cambridge, MA 02139, USA

## Abstract

Most animals coordinate behaviour using neural computations. Yet, single-celled organisms also exhibit stimulus-responsive, even cognitive, actions. To understand how a single cell can coordinate and drive complex behaviours without any neural encoding, we study an algal protist – a motile cell with four extremely long cilia. The organism displays a surprisingly rich locomotor repertoire, emerging from the intricate dynamics of the cilia, which form a tight bundle when swimming. We leverage high-speed quantitative live imaging to extract the spectrum of possible ciliary beating patterns, and derive a dispersion relation coupling the temporal frequency and spatial wavelength of cilia oscillations. We further reconstruct the attractor manifold embedded in the behavioural space, showing that despite the range and complexity of ciliary beating modes, the underlying behavioural manifold is intrinsically low-dimensional with elaborate topological structure. Dynamic and excitable transitions in motility behaviour are encoded as trajectories in this space.

---

Behaviour is shaped by an organism’s internal state and sensory experiences. Quantitative ethology has helped reveal the deep evolutionary origins of information processing and natural computation in behaving animals [1, 2]. In vertebrates, neuronal networks and complex patterns of electrical activity encode sensory inputs, control body posture, and coordinate movement gaits [3, 4]. In *C. elegans* crawling, the worm’s neuromuscular control architecture can be derived from dynamical systems representations of the worm’s posture changes [5, 6] In *Hydra*, a cnidarian with a simple distributed nervous system, ‘wireless’ signalling via neuropeptides controls sophisticated behaviours, such as somersaulting [7]. Yet, in the context of behaviour, life’s smallest organisms are often overlooked. Despite their miniscule size and relative simplicity, microorganisms routinely perform remarkably complex actions from directional sensing and tactic navigation to prey-capture, feeding, and even mating [8, 9]. All this is achieved without a nervous system or any musculature that we would typically find in animals executing similarly sophisticated tasks.

At the cell-scale, detailed study of behaviour is hindered by the challenge of imaging these processes at the requisite spatial and temporal resolution. Conventional brightfield microscopy cannot easily resolve subcellular features such as the characteristic shape changes of cilia or flagella, due to their fast dynamics, or intrinsically three-dimensional actuation patterns [10, 11]. For example, fluorescent labelling of flagella was needed to visualise how the enteric bacterium *C. jejuni* coordinates two opposing flagella for swimming by wrapping the leading flagellum around the cell body as it swims [12]. Despite advances in imaging and marker-less tracking [13], the quest to link the subcellular dynamics of microscopic appendages to whole-organism behaviour remains incomplete.

Here, we establish a motile marine alga (*Pterosperma*) as a powerful model system to investigate complex behavioural encoding in a single-celled organism. The species, which belongs to the basal lineage of Prasinophyte flagellates, is widely distributed and plays a critical role in marine ecosystems [14, 15]. During part of its life cycle, the cells form highly resistant cyst-like structures known as phycomata, which have even been observed in Silurian fossil records [16]. Each *Pterosperma* cell has a 10 *µ*m cell body and four extremely long cilia which emerge from an anterior groove (Fig. 1a). Its complex ciliary apparatus confers a unique form of fast motility, in which all four cilia tightly bundle to function as a single compound cilium [17]. The bundle maintains its integrity during forward swimming and even through diverse transitory manoeuvres. To capture the full behavioural space of this organism, we adopt a multi-scale approach spanning population dynamics down to single cilium activity. We leverage the organism’s highly planar ciliary beat waveform performing high-speed imaging to track and resolve ciliary shape dynamics over millisecond timescales. By decomposing these waveforms into welldefined shape modes and examining continuous transitions in behaviour, we reveal that the rich waveform dynamics are constrained by an empirically determined dispersion relation, and admits an intrinsically low-dimensional phenotypic space. This mirrors the low-dimensional nature of beat patterns found on reactivated ciliary axonemes (detached from the cell) [18].

**Figure 1.**
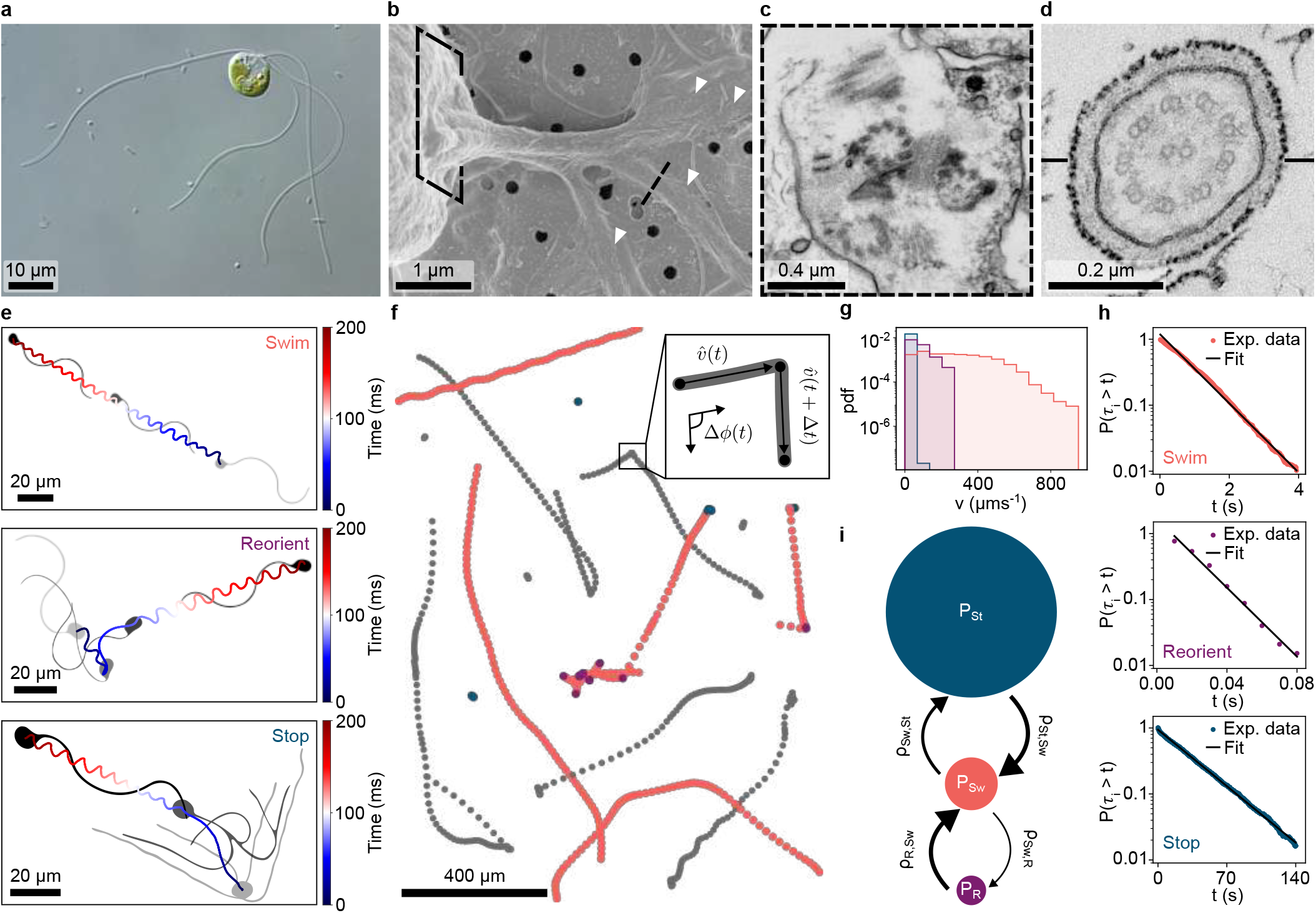
Organism morphology and free-swimming dynamics. **a**, *Pterosperma* cell imaged by differential interference contrast microscopy. **b**, Scanning electron microscopy image showing all four cilia emanating from the same location within the cell body, white arrows indicate the cilia. **c, d**, Transmission electron microscopy images (corresponding to the cross sections annotated by the black dashed lines in **b**) of the arrangement of basal bodies, striated fibres and ‘9 + 2’ ciliary axonemes. **e**, Illustrative cell body trajectories and ciliary arrangements during three distinct behavioural macrostates. From top to bottom: Swim, Reorient *→* Swim, Stop *→* Swim. **f**, Example free-swimming trajectories (see Supplementary Video 1), selected trajectories are coloured by state: Swim (coral), Reorient (purple), Stop (teal). Inset: schematic illustrating cell body velocity *v &* turning angle Δ*ϕ*. **g**, State specific cell speed histograms for n = 2096 trajectories. **h**, State survival probability *P* (*τ*_*i*_ > *t*) coloured circles indicate experimental values, exponential fits are shown with a solid black line. **i**, Transition network with statistics obtained from the Markov model. Steady state probabilities (*P*_Stop_, *P*_Swim_, *P*_Reorient_) = (0.966 *±* 0.002, 0.033 *±* 0.001, 0.00030 *±* 0.00005). Probability of each transition (*ρ*_St*→*Sw_, *ρ*_Sw*→*St_, *ρ*_Sw*→*R_, *ρ*_R*→*Sw_) = (1, 0.70 *±* 0.04, 0.30 *±* 0.04, 1).

Across multiple eukaryotic taxa, groups of cilia routinely self-assemble and organise into myriad specialised structures that mediate complex functions from sensing to motility, with minimal or no centralised control [19, 20]. Given the conserved nature of the cilium, our work uncovers general principles of how sophisticated locomotor gaits can arise from parsimonious actuation rules. In particular, our analysis reveals a vast functional spectrum of ciliary waveforms existing on the same structure, enriching our perspective on how dynein motors distributed along a cilium self-organise to produce emergent dynamics [21]. Taken together, we demonstrate how an aneural organism’s distinctive swimming behaviours are physically embodied in the spatiotemporal activity of the cilia actuators themselves.

## Quantification of free-swimming behaviour

The freely swimming *Pterosperma* cells studied here have a distinctive morphology consisting of an ellipsoidal cell body with dimensions (9 *±* 1) *×* (7.1 *±* 0.8) *µ*m, and four cilia that emerge from an anterior groove (Supplementary Fig. 1a). The cilia are unusually long (Fig. 1a), each with a typical length of 67 *±* 6 *µ*m (Supplementary Fig. 1b). Both body and cilia are covered with scales which form a primitive cell wall [14, 17, 22]. We visualised the ciliary apparatus using scanning and transmission electron microscopy to reveal that the cilia are anchored within the cell body by basal bodies that assume a 3 + 1 arrangement [17] (Fig. 1b). Fibrous structures connecting the basal bodies likely mediate cilia bundle synchrony, as shown in other microalgae [23] (Fig. 1c). All cilia have the highly conserved 9 + 2 axonemal structure (Fig. 1d).

We identified three behaviours: long periods in which the cell body remains stationary (Stop), followed by smooth forward swimming (Swim) interrupted by rapid reorientation events (Reorient). The organism typically unfurls its four cilia in the Stop state (Fig. 1e.), but always bundles them as a single ‘compound cilium’ during swimming and reorientations. Ciliary bundle formation is therefore temporary and arises from hydrodynamic/steric interactions; indeed, no physical connective components have been observed between the cilia except between the basal bodies [17]. In the Swim state, a travelling wave propagates down the cilium to drive forward propulsion while reorientations are achieved through large curvature bends of the compound cilium (Fig. 1e).

To reveal the divergent scales over which the three main behaviours are performed, we captured thousands of single-cell trajectories across the population using a 10× objective at 100 frames per second (fps) (Fig. 1f & Supplementary Video 1). Cells were imaged at 10 − 20 *µ*m above the glass substrate. Each state in our tripartite classification is automatically identified based on the instantaneous swimming speed *v*(*t*) and the cosine of the turning angle 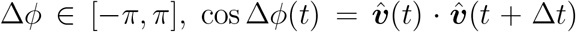 (Supplementary Fig. 2). The speed distribution (Fig. 1g) is broad, with *v* ≤ 20 *µ*m s^−1^ corresponding to Stop states. Reorientations are identified as epochs where cos Δ*ϕ <* 0.99 (Supplementary Fig. 2) and the swimming temporarily slows down.

This procedure automatically classifies the motility state as *i* ∈ {Stop, Swim, Reorient} across n = 2096 single-cell trajectories and 3947 transitions, to yield a robust quantification of the overall behaviour. State transitions can be modelled as a continuous time Markov chain (Supplementary section B.2), which admits the simple network structure shown in Fig. 1i. We calculated the mean state lifetimes ⟨*τ*_*i*_⟩, the steady state probability *P*_*i*_ of finding the cell in state *i*, the transition probabilities *ρ*_*ij*_ for the cell to switch from state *i* to state *j*, as well as the associated transition rate *q*_*ij*_. State survival probabilities *P* (*τ*_*i*_ > *t*) = *N* (*τ*_*i*_ > *t*)*/N* (*τ*_*i*_ > 0) (the probability for a cell to remain in a state for a time longer than *t*) [24, 25] decay approximately exponentially (Fig. 1h).

The three behavioural motifs occur over distinct timescales. Mean state lifetimes span three orders of magnitude in time from the highly persistent Stop state (⟨*τ*_*St*_⟩ = 58 *±* 2 s), through swimming (⟨*τ*_*Sw*_⟩ = 1.42 *±* 0.03 s), to the extremely short lived reorientations (⟨*τ*_*R*_⟩ = 0.041 *±* 0.002 s). Only specific transitions are allowable in this coarse-grained behavioural space: no Stop ↔ Reorient transitions are ever observed, therefore passage between these states must occur via the Swim state. We estimate the state probabilities *P* (*t →* ∞) = (*P*_Stop_, *P*_Swim_, *P*_Reorient_) = (0.966 *±* 0.002, 0.033 *±* 0.001, 0.00030 *±* 0.00005), so that the cell is extremely likely to be in the Stop state, possibly to conserve energy (see Discussion). We can also compute the probability of transitioning between each state (*ρ*_St*→*Sw_, *ρ*_Sw*→*St_, *ρ*_Sw*→*R_, *ρ*_R*→*Sw_) = (1, 0.70*±*0.04, 0.30*±*0.04, 1), noting that *ρ*_Sw*→*St_+ *ρ*_Sw*→*R_ = 1. The transition rates (*q*_St*→*Sw_, *q*_Sw*→*St_, *q*_Sw*→*R_, *q*_R*→*Sw_) = (0.0174*±*0.0005, 0.50*±* 0.01, 0.21 *±* 0.01, 24 *±* 1) s^−1^ fully characterise the motility dynamics.

## Active ciliary bundle oscillations drive distinct behaviours

To understand how these distinct behaviours emerge from the cell’s unique morphology and cilia dynamics, we performed high-magnification differential interference contrast imaging of single-cell swimming behaviour at 1000 fps using a 40*×* and 63*×* objective to capture ciliary bundle and single-cilium dynamics respectively. These high spatiotemporal resolution videos are used to resolve the intricate dynamics of the ciliary apparatus during each of the three behavioural macrostates (Supplementary Video 2).

Ciliary waveforms were extracted from each frame using custom MATLAB^®^ code using the medial axis transform [26] to delineate between the cell body and cilium proper. We focus on the dynamics of the centreline of the ciliary bundle, fully described by its instantaneous tangent angle *θ*(*s, t*) parametrised by arc length *s* and time *t* (Fig.2a., Supplementary section C). Distinct ciliary beating patterns underlie the highly variable cell propulsion modes observed in Fig. 2d.

**Figure 2.**
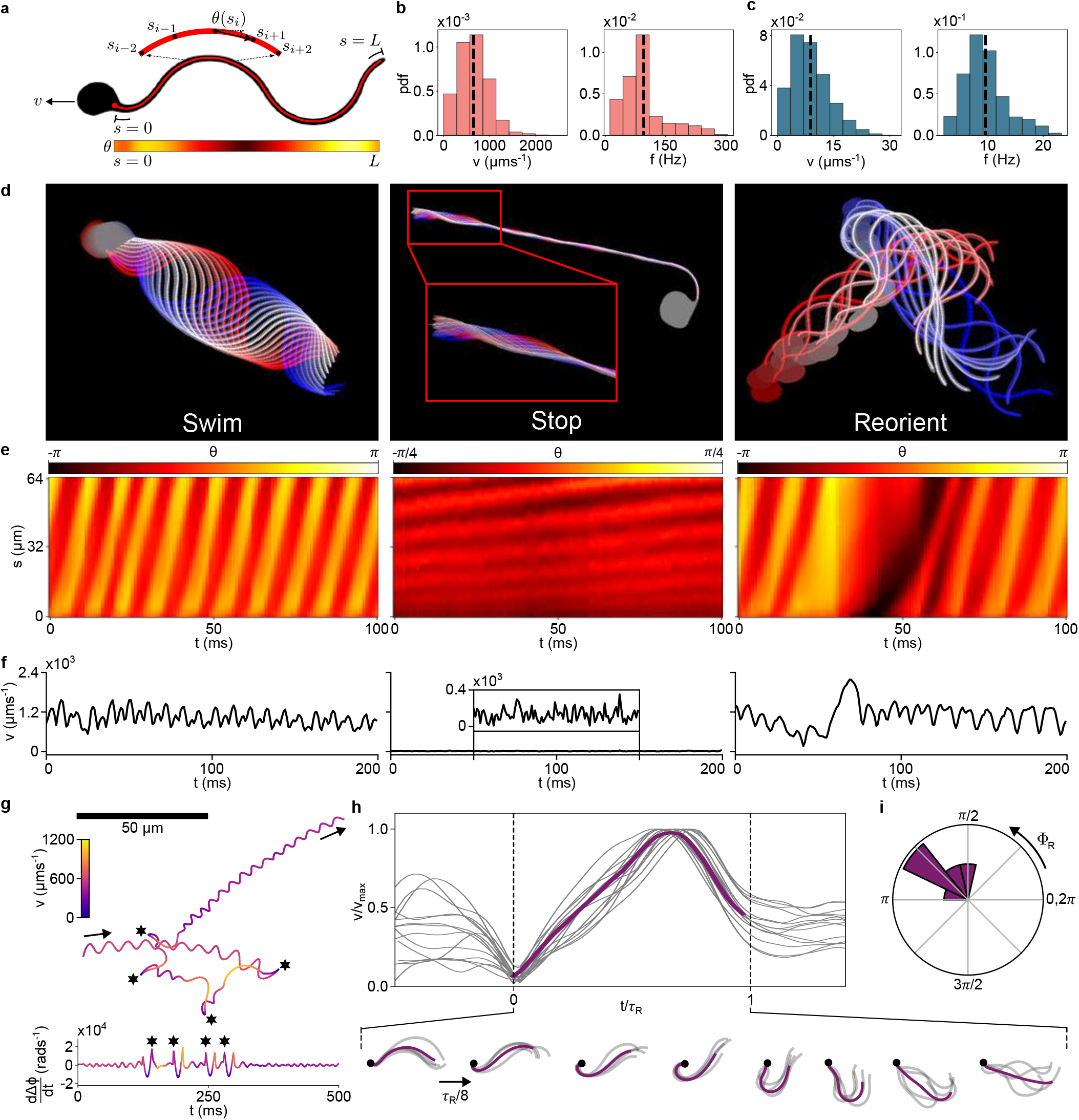
Ciliary bundle displays complex waveforms. **a**, Ciliary (bundle) waveform parametrised by tangent angle and arc length *θ*(*s*). **b**-**c**, Histograms of cell body speed (left) and ciliary beat frequency (right) during Swim (n_cells_ = 92) and Stop (n_cells_ = 14, n_cilia_ = 16) states. Dashed black line indicates mean value. ⟨*v*_Swim_⟩ = 646 *±* 326 *µ*m s^−1^, ⟨*f*_Swim_⟩ = 95 *±* 53 Hz and ⟨*v*_Stop_⟩ = 9 *±* 5 *µ*m s^−1^, ⟨*f*_Stop_⟩ = 10 *±* 4 Hz (mean *±* s.d.) **d**, Cell silhouette with cilium trace overlaid and colour coded by time, for each behavioural macrostate. **e**, Kymographs of *θ*(*s, t*) (arc length *s* and time *t*) corresponding to each behaviour above. **f**, Sample timeseries of cell speed *v*(*t*) in each state. **g**, Trajectory of cell undergoing sequential reorientations (top) with corresponding angular speed profile (bottom). Asterisks indicate reorientations. **h**, Cell speed profiles, normalised by the max speed *v*_*max*_ and reorientation time *τ*_*R*_, show reorientation dynamics are stereotyped (n_cells_ = 11, n_reorientations_ = 14). Inset: Phase-ordered waveforms traced during reorientations (thick lines: waveforms averaged across 4 cells). **i**, Circular histogram of reorientation angle Φ_*R*_ (n_cells_ = 11, n_reorientations_ = 14).

In the Swim state, the cilia remain bundled. Robust base-to-tip travelling waves propagate with highly variable frequency and amplitude. The distribution of frequencies ranges from 12 to 304 Hz with a mean of 95 Hz, resulting in swimming speeds varying across two orders of magnitude (Fig. 2b.) (n_cells_ = 92). Such a broad spectrum contrasts with previous measurements from other organisms and sperm flagella, where cilia beat frequencies are typically more tightly defined for any given species [27, 28, 29, 30].

The Stop state is associated with minimal body movement yet active cilia oscillations (Fig. 2c.). Here, robust base-to-tip directed waves are observable on the distal 3*/*4 of the cilium, with mean beat frequencies of 9.5 ± 0.1 Hz (Fig. 2c, d) (n_cells_ = 14, n_cilia_ = 16). However, due to the randomised orientation of the unbundled cilia, the cell motion resulting from these cilia oscillations is largely non-progressive, with average cell migration speed 8.83± 0.01 *µ*m s^−1^ and diffusivity of only 0.2 *µ*m^2^ s^−1^ (Supplementary Section B.3). The functional role of this novel oscillatory mode of motile cilia remains unclear but may be to enable active sensing [24].

Finally, the Reorient mode is detected as an abrupt change in the swimming trajectory and speed (Fig. 2g). This results from rapid changes in the ciliary bundle waveform (which remains intact), in contrast to the quasi-steady oscillations displayed by the cilia during the Swim and Stop states (Fig. 2d-f). Aligning 14 reorientations from 11 cells we found them to be highly stereotyped (Supplementary Fig. 5). Typically, a large initial bend develops at the cilium base which rapidly brings the cell to a halt, rotating the cell body through ∼ 180°. This bend propagates down the cilium propelling the cell rapidly forward, followed by a relaxation back into the Swim state, during which waveform stereotypy temporarily breaks down. We estimate that reorientations produce a stereotyped turning angle peaked around 130± 30° with a duration of 36 ± 3 ms (Fig. 2i & Supplementary Fig. 5). Reorientations are an essential component of the organism’s tripartite navigation strategy, akin to ‘tumbles’ in other microswimmers [31, 32].

### High-resolution spectral decomposition of ciliary beating modes

So far, we have established that *Pterosperma* displays many distinct behaviours driven by stereotyped dynamics of the cilia. To reveal the temporal evolution of different ciliary oscillation patterns, we seek a unifying framework capable of capturing this extraordinary complexity using a common shape basis for all observed beat waveforms. Methods previously employed to capture shape information from undulatory locomotion based on PCA or Fourier modes are either nondifferentiable or cannot adequately resolve the highly transient, non-oscillatory modes we observe during reorientations. Instead, we decompose ciliary bundle shapes using Chebyshev polynomials of the first kind *T*_*n*_(*s*) [33]. Such a basis is advantageous since Chebyshev expansions converge spectrally for non-periodic smooth functions allowing for accurate shape reconstruction [34]. They also admit analytical derivatives and the coefficients are interpretable physically as bending mode amplitudes.

We express the instantaneous shape of the cilium (or ciliary bundle) as a linear superposition of Chebyshev modes (Fig. 3a.) weighted by their respective time-dependent amplitudes 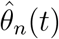:

**Figure 3.**
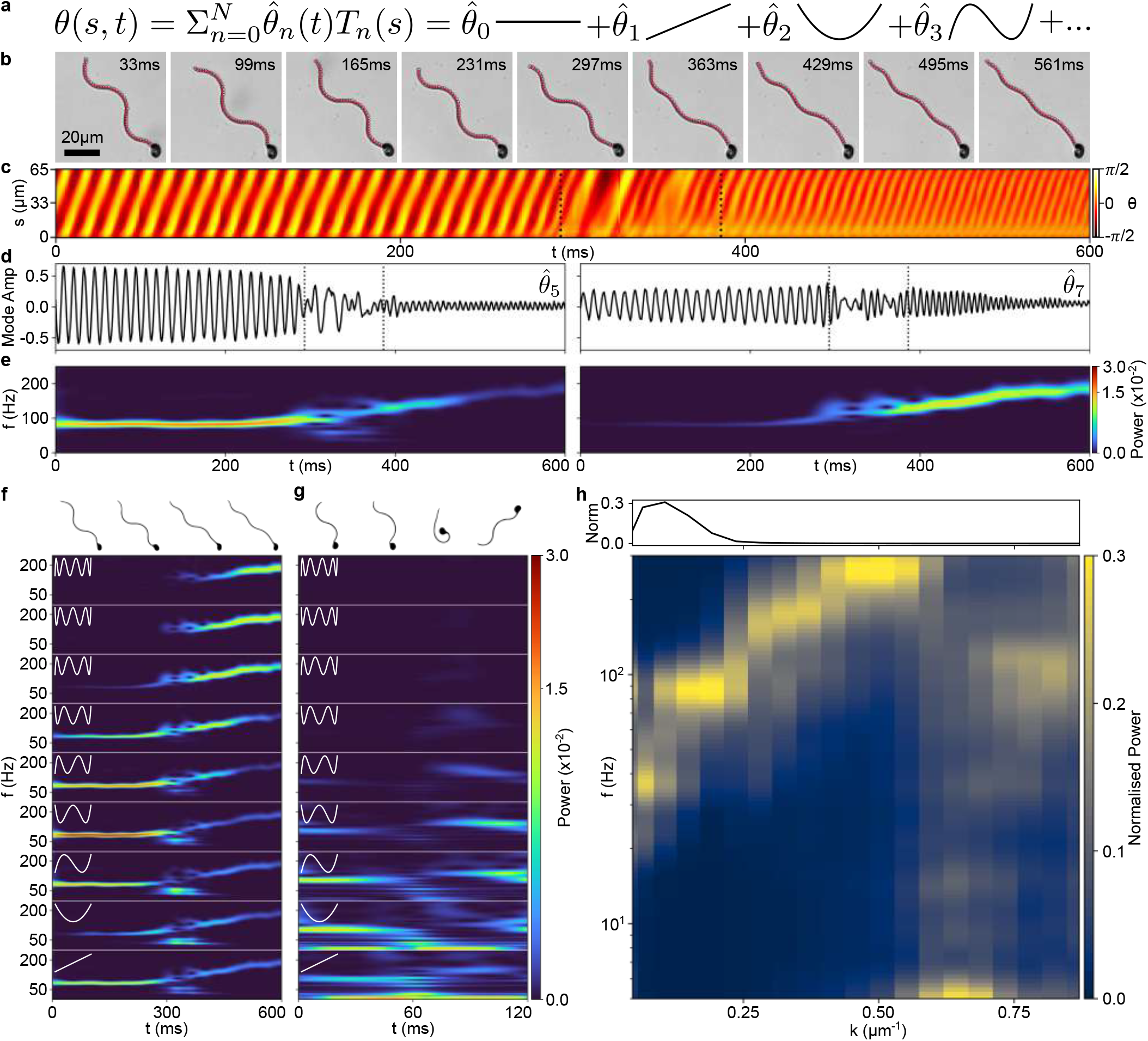
Spectral decomposition of ciliary bundle dynamics. **a**, Pictorial representation of the instantaneous ciliary bundle shape as a superposition of Chebyshev modes. **b**, Raw video frames of a mode transition sequence overlaid with the Chebyshev waveforms reconstructed with 20 modes. **c**, Kymographs of tangent angle representation *θ*(*s, t*) (same dataset as **b**). **d**,**e**, Selected Chebyshev mode amplitudes 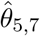 corresponding to the ciliary bundle traces in **b**, and their normalised continuous wavelet transforms, **Ŵ** _Ψ_(*f, t*). **f, g**, Wavelet vectors, modes 1 − 9, for a swimming cell modulating its beat pattern (**f**, same sequence in **b**-**e**) and a cell performing a reorientation (**g**). **h**, Dispersion relation f-k for the entire dataset of n = 219, 368 ciliary bundle waveforms, from 125 cells. The magnitude at each k (top) is used to normalise the heatmap.

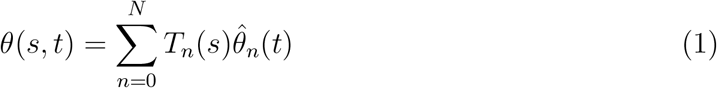

where the coefficients are determined by first fitting an expansion in Chebyshev polynomials along (*x, y*) to determine 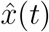 and *ŷ*(*t*) using a trust region regularised least squares, motivated by the convergence properties of Chebyshev polynomials for smooth functions. Using the Chebyshev expansions of (*x, y*) the curve can then be efficiently re-parametrised to an arclength representation *θ*(*s, t*) determined using *θ*(*s, t*) = arctan(*∂*_*s*_*y*(*s, t*)*/∂*_*s*_*x*(*s, t*)) where the derivatives are calculated from the spectral representation. The theta coefficients are then obtained using Chebyshev projection on *θ*(*s, t*). The 0^th^ coefficient of the decomposition corresponds to a weighted instantaneous orientation of the cilium with respect to the lab frame horizontal axis (Supplementary Fig. 4d) and the 1^st^ coefficient to a weighted average curvature, while higher modes describe more detailed cilium shape features. Here we used a total of *N* = 20 modes to capture the entire ciliary shape space with a root mean square reconstruction error of 0.368 *µ*m on average (Supplementary Fig. 4c).

We apply this decomposition to high-resolution sequences of *Pterosperma* cilia. To capture both the dominant shape modes at each time point as well as its time-varying frequency content (Fig. 3b-e), we compute the continuous Morlet wavelet transform 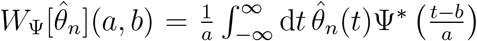 of the mode amplitudes [35], for different discrete values of the wavelet frequency scale 1*/a*_*l*_ and time shift *b*_*k*_ = *t*_*k*_, resulting in a wavelet matrix 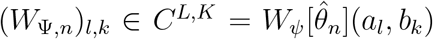. For every observed ciliary waveform, we can ‘stack’ the wavelets for each mode of the decomposition into a unique ‘wavelet vector’ :

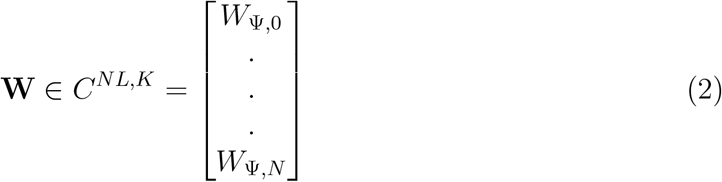

to reveal the wavenumber- and time-dependent frequency information embedded within these complex waveforms. Since the magnitude of the wavelet transform is proportional to the amplitude of an oscillation, we chose to normalise each column of the wavelet vector to have unit norm **Ŵ** (*t*_*k*_) = **W**(*t*_*k*_)*/* | **W**(*t*_*k*_) |, to ensure good resolution across cilia states of varying amplitudes while maintaining the relative importance of each Chebyshev mode within a given state.

We illustrate the results with two very different behaviours. In the first example, a swimming cell is actively modulating the dynamics of its ciliary bundle (Fig. 3b,c). The corresponding wavelet scalograms are plotted for each Chebyshev mode, where |*W*_Ψ_(*f, t*) | ^2^ represents the signal power at each frequency over time (Fig. 3d, e). The time-evolution of wavelet amplitudes reveals unprecedented spectral structure within the swimming state itself (Fig. 3f), allowing us to visualise the rapid but continuous modulation of the large beat amplitude 80 Hz wave to a smaller-amplitude 180 Hz wave. The gradual decay of Chebyshev modes 1 − 5 coincides with the appearance of signal in modes 7− 9 concurrent with an increasing beat frequency (Fig. 3f). The second example shows a cell undergoing a fast reorientation, whose ciliary dynamics are captured similarly (Fig. 3g). An initially well-defined oscillation is scrambled, with the dominant wavelet amplitudes shifting to the lowest Chebyshev modes. As the cell relaxes back to a new swimming state, the intensity signal percolates from the higher to lower modes before the cell settles into its swimming state again now dominated by a new, slightly higher, beat frequency.

We apply this spectral-wavelet analysis to a large high-resolution dataset of single-cell ciliary dynamics consisting of 219, 368 frames of ciliary bundle shapes from a total of 125 cells. From the instantaneous dynamics of the cilium or ciliary bundle we can extract dominant temporal frequencies *f* corresponding to a given spatial frequency or wavenumber *k* = 2*π/λ* by time-averaging *W*_Ψ,*n*_ across cells and assigning to each Chebyshev mode its dominant wavenumber (Fig. 3h).

Our comprehensive dataset captures a remarkable dynamic range in both temporal and spatial frequencies, and occupies two distinct branches in *f* -*k*-space, revealing a ‘dispersion relation’ that constrains ciliary beating. The prominent diagonal band from low to high *f* corresponds to waveforms observed during swimming; cilia transitioning between different beating modes within the Swim macrostate undergo bidirectional movement or fluctuations along this branch (Supplementary Video 3). The swimming dynamics obey an approximately linear dispersion relation with *f* ∼ *k* (Supplementary Fig. 6a). The quantisation of the oscillations into four prominent frequency bands, occurring at 37, 88, 184, and 265 Hz (Supplementary Fig. 6b), is a further hallmark of the underlying dynein-driven beat generation mechanism (see discussion below). Meanwhile, the vertical band at low *f* and high *k* corresponds to transitions from the swimming macrostate into the Stop state, usually accompanied by a dramatic decrease in beat frequency (Supplementary Video 3). Exit from a Stop state is driven by a discontinuous transition in beat mode from a low-*f*, high-*k* state back onto the main diagonal branch.

These sub-cellular dynamics reveal the existence of novel cilia-driven sub-states in the motility repertoire of *Pterosperma* that are completely inaccessible from cell body tracking alone. This also explains the extremely broad distributions of beat frequency and cell speed we measured previously, which do not arise from population heterogeneity but from temporal variability in single cell behaviour.

### Latent manifold captures whole-cell behavioural transitions

We proceed to reconstruct the behavioural state space underlying the entire spectrum of ciliary dynamics a single *Pterosperma* cell can perform.

To understand the spatial partitioning of behaviour according to the different modes in Fig. 3h, we identified nine representative waveform sequences and plotted them alongside their corresponding dispersion relations in Fig. 4a. The entire space of beat patterns associated with whole-cell behavioural transitions can be visualised as a single manifold reconstructed from a small number of dimensions (Fig. 4b). This is obtained by performing a non-linear dimensionality reduction via multidimensional scaling (MDS) on a distance matrix calculated from the normalised stacked wavelet amplitudes *d*_*ij*_ = |**Ŵ** (*t*_*i*_) − **Ŵ** (*t*_*j*_)|. Just three MDS dimensions were sufficient to capture 83% of the variance in the data, while six dimensions captures over 95% (Supplementary Fig. 7).

**Figure 4.**
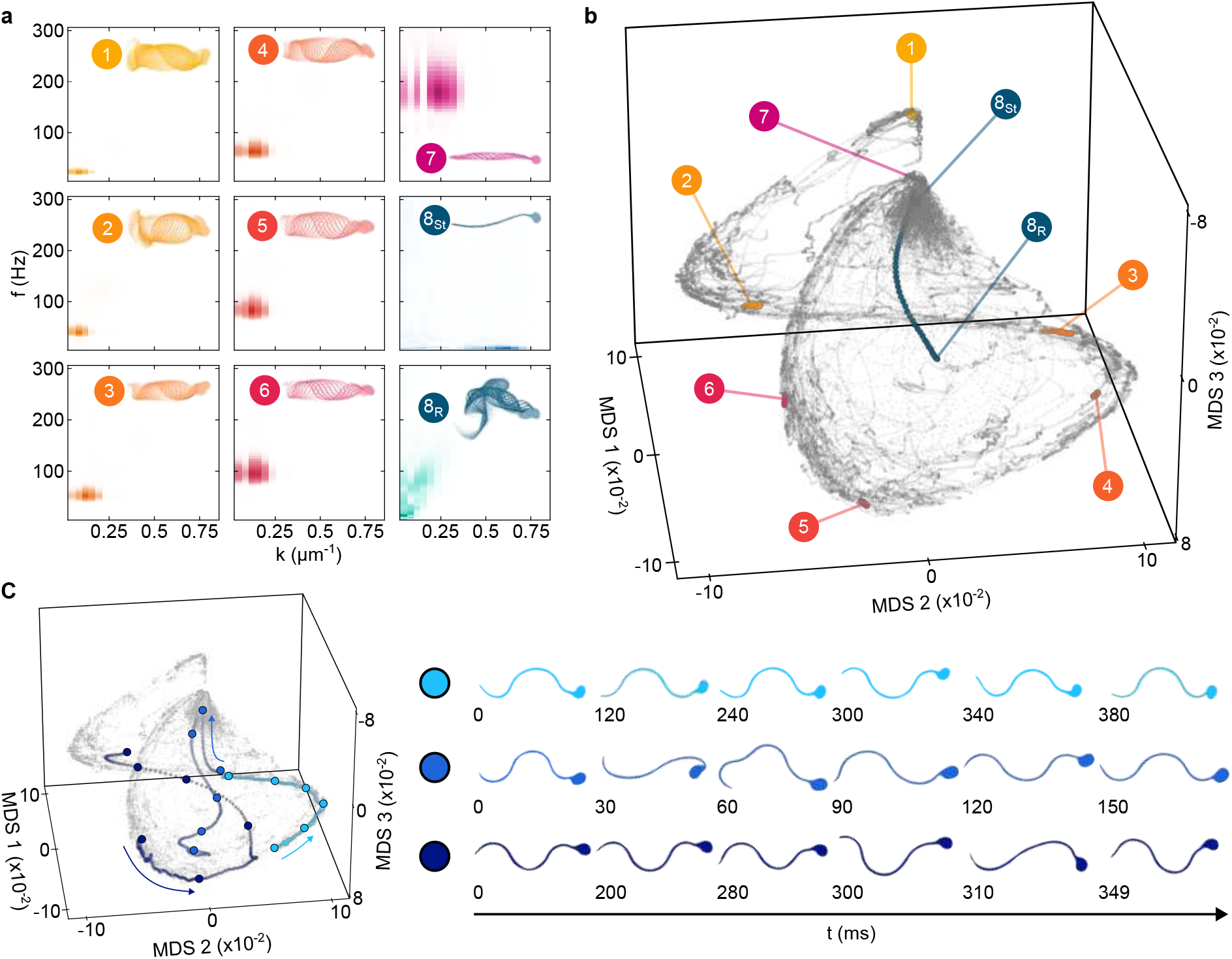
Excitable dynamics lies on a low-dimensional behavioural manifold. **a**, Normalised dispersion plots computed for nine illustrative waveform sequences. **b**, *Pterosperma’s* behavioural manifold illustrated in the first three MDS dimensions. Grey points indicate the positions on the manifold of all n = 219, 368 ciliary bundle waveforms. The manifold coordinates of the waveform sequences in **a** are indicated. **c** Three behavioural trajectories on the manifold. Large filled circles indicate the positions of the cell snapshots (right) and arrows the direction of time.

Illustrated here in three MDS dimensions with the nine waveform sequences from Fig. 4a highlighted, the behavioural manifold is two-lobed and highly structured. Following the dynamics along the manifold, we see that beat patterns associated with Swim states define an elaborate envelope bounding all other states. Transient behaviours appear as excursions from the main envelope, where the trajectory ventures into the space off the envelope and then rejoins it far from where they left resulting in a sharp change in the steady swimming behaviour (Supplementary Video 4).

For the most part, trajectories flow bidirectionally, following an inverse correlation between beat amplitude and frequency. Yet, nearby phase trajectories can diverge substantially over time, a clear signature of chaotic dynamics (Fig. 4c). In contrast, certain behaviours produce trajectories that only flow in one direction and are therefore irreversible. Reorientations travel into the cluster containing the Stop state tending to positive values of MDS-1, where the cilia exhibit large deformations. Stop and reorientation events co-localise as each correspond to very low frequency modes (8_St_ and 8_R_). Meanwhile, cells transitioning towards their Stop state from swimming typically do so by ramping up the oscillation frequency and reducing the amplitude. Such trajectories cross the manifold peak from the negative MDS-1 side and reach a point of no return, beyond which cells are committed to entry into their Stop state. Exiting the Stop state involves a rapid excursion out of the Stop cluster (excitatory phase) back onto the swimming envelope followed by a refractory period.

### Geometric encoding of embodied behaviour

Behavioural studies of microorganisms often target a single scale, either by capturing high-resolution appendage dynamics from immobilised cells [36, 27, 37], or using low-resolution centroid-based tracking to follow whole-organism movement [31, 32, 38]. The latter coarse-graining may overlook an organism’s nuanced posture or shape dynamics [5, 2]. Here, we developed a multiscale approach to reconstruct the state-space underlying all natural behaviours of a single microswimmer, focussing on the oscillatory dynamics of its cilia. This representation exposes the existence of continuous manifold structure through a non-linear embedding of high-dimensional timeseries. In this space, two behaviours are considered similiar if they correspond to neighbouring points, resolving stereotyped dynamics by distance similiarity. By mapping behavioural dynamics to trajectories on a low-dimensional attracting manifold, we capture not only the geometric encoding of the states themselves but also the relationships between different states.

For a ciliated single cell, whose motility is dominated by geometric mechanics at zero-Reynolds number [39, 40], there is a clear equivalence between the intrinsic shape dynamics of the cilia and whole-cell behaviour. Key features of this embodied dynamical system must therefore derive from the underlying ciliary actuation mechanism. Here in live, freely behaving *Pterosperma* cells, the rich motility dynamics of its ciliary bundle collapse onto a low-dimensional manifold with striking, well-defined structure. This mirrors the intrinsic low-dimensionality of isolated *Chlamydomonas* cilia that have been detached from the cell body and reactivated with ATP under different conditions [18].

From the spectrum of *Pterosperma* ciliary dynamics we also uncovered an empirical dispersion relation [41, 42]: cilia cannot oscillate arbitrarily but are constrained by certain allowable temporal frequencies *ω* = 2*πf* and spatial wavenumber *k*. This has profound consequences for waveform generation by dynein co-operativity in cilia and (eukaryotic) flagella, which remains incompletely understood [21, 43, 44, 45, 46]. For an active elastic filament actuated by internal sliding forces, a linearised force balance suggest a complex dispersion relation of the form *ωξ*_*⊥*_*i* = −(*EI*)*k*^4^ + *a*^2^*iωχ*(*ω*)*k*^2^, where *EI* is the filament’s bending rigidity, *a* the centre-to-centre separation, and *ξ*_*⊥*_ the perpendicular drag coefficient. Under certain assumptions of motor kinetics *χ*(*ω*), this admits spontaneous oscillations and travelling waves [47]. This type of self-organised dynamics could produce the discretisation of beat modes we found in Fig. 3h, as also observed in *Chlamydomonas* cilia undergoing phase slips [27]. Future models of ciliary actuation should reproduce temporal sequences of ciliary beat patterns, not just isolated snapshots of their steady-state dynamics.

The various behaviours exhibited clear timescale separation, from fast reorientations (*<* 50 ms), to intermediate timescales of swimming (∼ 1 s), to the long-lived Stop states (> 50 s). Our geometric embedding provides new insights into the mechanism of reversible shape transitions over short times, and the irreversible dynamics of entry and exit from Stop states over long times. The small-amplitude oscillations exhibited by the cilia during the quiescent state suggests that the system operates close to criticality [48, 24]. By analogy with hair-cell oscillations [49], dynamics near a limit cycle rather than fixed point may enable fast responses, corresponding to rapid excursions on our behavioural manifold (Fig. 4b). In an ecological context, ciliary mechanosensitivity likely enables planktonic algal flagellates to respond rapidly to predation or other sources of disturbance [50, 51, 52]. Signatures of dynamical excitability may have been a critical cellular innovation [9].

Behaviour exposes the nature of sensorimotor representations in organisms. In a soft body, the interplay between deformable morphology and embodied behaviour can effectively orchestrate complex dynamics based on simple actuation rules, applicable to shape-programmable materials [53, 54]. In excitable cells whose motility is shaped by the intrinsic oscillations of the cilia, a single control parameter such as Ca^2+^, cAMP, or ATP concentration, could nonlinearly select, for example via Hopf bifurcation, a wide dynamic range of physical parameters such as beat frequency or waveform without explicit control of dynein function. Our analysis presents a new framework for exploring the emergence of complexity in single-celled organisms.

## Methods

### Cell Culture

*Pterosperma sp*. (K-1082, Norwegian Culture Collection of Algae) was cultured by suspending 3 mL of 1-week old cell culture in 30 mL of TL30 media (https://norcca.scrol.net/node/3956). Cells were grown at 21 °C and 40 % humidity on a 14 : 10 hour light : dark cycle at a light intensity of 2, 000 lux. Cultures were harvested for experiments 4 − 6 days after inoculation during the late exponential phase of growth at concentrations of ∼ 10^5^ cells*/*mL.

### Electron microscopy

For electron microscopy studies cells were fixed in 2% glutaraldehyde and 2% paraformaldehyde in 0.1M sodium cacodylate buffer (pH 7.2) for 1 hr at room temperature. Fixed cells were washed 3*×* for 5 min in the same buffer and post-fixed for 1 hr in 1% osmium tetroxide.

For Transmission Electron Microscopy (TEM), following [55], the post-fixed sample was additionally reduced in 1.5% potassium ferrocyanide in 0.1 M sodium cacodylate buffer.

In both Scanning Electron Microscopy (SEM) & TEM cells were washed three times for 5 min each in deionised water before dehydration through a graded ethanol series.

SEM samples were passed through a 200nm polycarbonate filter using a mild vacuum. After a 3 min incubation in hexamethyldisilizane [HMDS, Merck, Gillingham, UK], filters were air-dried and mounted onto aluminium stubs with double-sided sticky carbon tabs. Samples were sputter-coated with a 10nm layer of gold/palladium and imaged using a Zeiss Gemini 500 scanning electron microscope.

TEM samples were gradually infiltrated with Durcupan resin [Sigma Aldrich, Merck, Gillingham, UK] and embedded in BEEM capsules. The resin was polymerised overnight at 60°C. 70nm sections were collected on pioloform-coated EM copper grids [Agar Scientific, Rotherham, UK] and post-contrasted with Reynold’s lead citrate. Imaging was performed using a JEOL JEM 1400 transmission electron microscope operated at 120 kV, with images captured using a Gatan ES1000W digital camera.

### Brightfield microscopy

Live cell imaging chambers were constructed by spin coating ∼ 0.5mm Polydimethylsiloxane (PDMS) sheets, punching 6mm diameter circular wells and securing this section between two cover glass slides.

Imaging was performed 10 − 20 *µ*m above the surface using an inverted microscope [Leica Microsystems DMi8] with a high speed camera [Phantom Vision Research V1212]. Videos for population analysis were taken under brightfield (BF) illumination at 100 fps using a 10× (HC PL/0.40) dry objective. For higher-resolution analysis, differential interference contrast (DIC) images were obtained at 1000 fps using a 40 × (HC PL/0.6) dry objective for swimming and reorientation single cell analysis and a 63 × (HC PL/1.2) water immersion objective for cells in their Stop state with a DIC prism providing pixel resolutions of 0.370 *µ*m*/*pixel and 0.234 *µ*m*/*pixel respectively.

### Cilium Tracking and Reconstruction

A custom cilium tracking protocol was implemented in MATLAB^®^ to extract pixel-wise traces from videos of cell dynamics. Single cell videos are binarised, and the cilium centre line extracted using a threshold on the medial axis transform (MAT) [26] (Supplementary Fig. 4a).

We approach the parametrisation of the pixel-wise traced ciliary waveform using a shape space decomposition. Here we leverage Chebyshev polynomials of the first kind *T*_*n*_(*s*). These are selected because they are analytically differentiable and effective at capturing non-periodic functions, such as those found in the broad spectrum of ciliary shapes performed by *Pterosperma*, with a small number of modes. Here we provide an overview of the technique, full mathematical details are provided in the supplementary information.

The raw pixel-wise traces **r**(*s, t*) = [*x*(*s, t*), *y*(*s, t*)] can be described by the linear superposition of Chebyshev modes weighted by their time dependent amplitudes 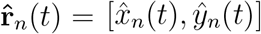 for *n*∈ {0, 1 … *N*}. The mode amplitudes are obtained with a trust region regularised least squares fit. This requires the constraint that the coefficients should decay exponentially under the assumption that our waveforms are smooth functions [34] and a regularisation term on the total variation of the second derivative which enforces smoothness by penalising large changes in the second derivative. In this step *N* = 125 coefficients are fitted providing a root mean square error in the reconstruction of less than the size of a pixel on average. Not all coefficients are necessary to describe the shape but are important for calculating smooth, high quality derivatives important for re-parametrisation and computing the angular representation.

From the Cartesian space curve correctly re-parametrised by the arc length a smooth transformation to the tangent angle representation can be achieved:

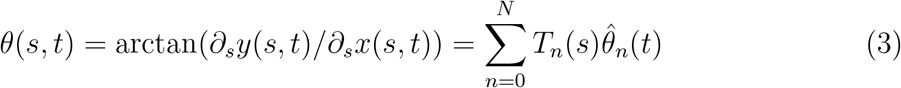

where the subscript *s* indicates a derivative with respect to arc length.

The Cartesian space representation can be recovered using Equation 4 (Supplementary Fig. 4b):

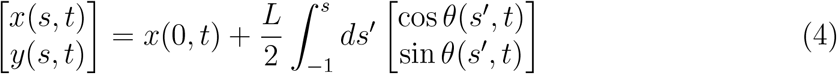

where *L* is the total trace length of the cilium. The tangent angle Chebyshev coefficients, 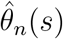, are defined by Equation 5:

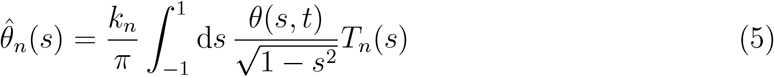

where *k*_0_ = 2 and *k*_*n*_ = 1 otherwise. The 0^*th*^ order coefficient corresponds to a weighted average orientation (Supplementary Fig. 4d) while the 1^*st*^ order coefficient represents a weighted average of the signed curvature.

For each Chebyshev polynomial an approximate wavelength can be assigned to the *n*^*th*^ polynomial as 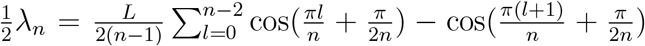 which is the mean distance between the extrema scaled by the average cilium length. The factor of 2 in the length scaling is due to the natural interval for Chebyshev polynomials being [−1, 1].

### Wavelet analysis and multidimensional scaling analysis

The wavelet transform is performed via the fast Fourier transform using the Morlet wavelet with characteristic frequency 2*π*. The initial and final 20 frames are cropped from the resulting transform to remove boundary artifacts. The transform is performed for each Chebyshev mode trajectory separately.

The dispersion relation is calculated by averaging the wavelet transform for each Chebyshev mode across all time points and chosen experiments. This averaging results in a single vector as a function of frequency. A dispersion heatmap is formed by stacking each time averaged wavelet vector across Chebyshev mode.

To calculate the multidimensional scaling (MDS) between each instantaneous stacked wavelet vector the data are randomly distributed into three bins with 1666 time points the same between the bins. Classical MDS is performed separately on each bin using Euclidean distances between the wavelet vectors. Since MDS is only defined up to an affine transformation, the resulting MDS coordinates are then aligned by fitting an affine transform between the overlapping points in each bin to produce a combined MDS embedding [56].

## Data Availability

All datasets necessary to reproduce the work detailed in this manuscript will be deposited at [URL to be inserted after review].

## Code Availability

Custom tracking and analysis codes associated with this manuscript will be deposited at [URL to be inserted after review].

## Acknowledgments

We would like to thank C. Hacker & P. Cherek for preparation of samples for SEM and TEM imaging, and R. Chait for stimulating discussions. This work was funded by the European Research Council (ERC) under the European Union’s Horizon 2020 research and innovation programme grant 853560 Evomotion (to KYW) and the NSF-Simons National Institute for Theory and Mathematics in Biology (NITMB) Fellowship supported via grants from the NSF (DMS-2235451) and Simons Foundation (MPS-NITMB-00005320) (to ADH). This research received support through Schmidt Sciences, LLC (Polymath Award to JD) and the MIT MathWorks Professorship Fund (JD).

## Ethics declarations

None

## References

[1] Pereira, T. D., Shaevitz, J. W. & Murthy, M. Quantifying behavior to understand the brain. Nature Neuroscience 23, 1537–1549 (2020).

[2] Brown, A. E. & De Bivort, B. Ethology as a physical science. Nature Physics 14, 653–657 (2018).

[3] Goulding, M. Circuits controlling vertebrate locomotion: moving in a new direction. Nature Reviews Neuroscience 10, 507–518 (2009).

[4] Kiehn, O. & Dougherty, K. Locomotion: Circuits and Physiology, 1623–1651 (Springer, Cham, Switzerland, 2022).

[5] Ahamed, T., Costa, A. C. & Stephens, G. J. Capturing the continuous complexity of behaviour in Caenorhabditis elegans. Nature Physics 17, 275–283 (2021).

[6] Pierce-Shimomura, J. T. et al. Genetic analysis of crawling and swimming locomotory patterns in C. elegans. Proceedings of the National Academy of Sciences 105, 20982–20987 (2008).

[7] Yamamoto, W. & Yuste, R. Peptide-driven control of somersaulting in Hydra vulgaris. Current Biology 33, 1893–1905 (2023).

[8] Coyle, S. M., Flaum, E. M., Li, H., Krishnamurthy, D. & Prakash, M. Coupled active systems encode an emergent hunting behavior in the unicellular predator Lacrymaria olor. Current Biology 29, 3838–3850 (2019).

[9] Wan, K. Y. & Jékely, G. Origins of eukaryotic excitability. Philosophical Transactions of the Royal Society B 376, 20190758 (2021).

[10] Rossi, M., Cicconofri, G., Beran, A., Noselli, G. & DeSimone, A. Kinematics of flagellar swimming in Euglena gracilis: Helical trajectories and flagellar shapes. Proceedings of the National Academy of Sciences 114, 13085–13090 (2017).

[11] Striegler, M., Diez, S., Friedrich, B. M. & Geyer, V. F. Twist–torsion coupling in beating axonemes. Nature Physics 21, 599–607 (2025).

[12] Cohen, E. J. et al. Campylobacter jejuni motility integrates specialized cell shape, flagellar filament, and motor, to coordinate action of its opposed flagella. PLoS pathogens 16, 1–24 (2020).

[13] Mathis, M. W. & Mathis, A. Deep learning tools for the measurement of animal behavior in neuroscience. Current opinion in neurobiology 60, 1–11 (2020).

[14] Parke, M., Boalch, G., Jowett, R. & Harbour, D. The genus Pterosperma (Prasinophyceae): species with a single equatorial ala. Journal of the Marine Biological Association of the United Kingdom 58, 239–276 (1978).

[15] Maruyama, S. & Kim, E. A modern descendant of early green algal phagotrophs. Current Biology 23, 1081–1084 (2013).

[16] Colbath, G. K. Fossil prasinophycean phycomata (Chlorophyta) from the Silurian Bainbridge Formation, Missouri, USA. Phycologia 22, 249–265 (1983).

[17] Inouye, I., Hori, T. & Chihara, M. Absolute configuration analysis of the flagellar apparatus of Pterosperma cristatum (Prasinophyceae) and consideration of its phylogenetic position. Journal of Phycology 26, 329–344 (1990).

[18] Geyer, V. F., Howard, J. & Sartori, P. Ciliary beating patterns map onto a lowdimensional behavioural space. Nature Physics 18, 332–337 (2022).

[19] Gilpin, W., Bull, M. S. & Prakash, M. The multiscale physics of cilia and flagella. Nature Reviews Physics 2, 74–88 (2020).

[20] Laeverenz-Schlogelhofer, H. & Wan, K. Y. Bioelectric control of locomotor gaits in the walking ciliate euplotes. Current Biology 34, 697–709 (2024).

[21] Howard, J., Chasteen, A., Ouyang, X., Geyer, V. F. & Sartori, P. Predicting the locations of force-generating dyneins in beating cilia and flagella. Frontiers in Cell and Developmental Biology 10, 995847 (2022).

[22] Becker, B., Marin, B. & Melkonian, M. Structure, composition, and biogenesis of prasinophyte cell coverings. Protoplasma 181, 233–244 (1994).

[23] Wan, K. Y. & Goldstein, R. E. Coordinated beating of algal flagella is mediated by basal coupling. Proceedings of the National Academy of Sciences 113, E2784–E2793 (2016).

[24] Wan, K. Y. & Goldstein, R. E. Time irreversibility and criticality in the motility of a flagellate microorganism. Physical Review Letters 121, 058103 (2018).

[25] Perez Ipiña, E., Otte, S., Pontier-Bres, R., Czerucka, D. & Peruani, F. Bacteria display optimal transport near surfaces. Nature Physics 15, 610–615 (2019).

[26] Walker, B. J., Ishimoto, K. & Wheeler, R. J. Automated identification of flagella from videomicroscopy via the medial axis transform. Scientific Reports 9, 5015 (2019).

[27] Wan, K. Y., Leptos, K. C. & Goldstein, R. E. Lag, lock, sync, slip: the many ‘phases’ of coupled flagella. Journal of the Royal Society Interface 11, 20131160 (2014).

[28] Ma, R., Klindt, G. S., Riedel-Kruse, I. H., Jülicher, F. & Friedrich, B. M. Active phase and amplitude fluctuations of flagellar beating. Physical Review Letters 113, 048101 (2014).

[29] Bayly, P., Lewis, B., Kemp, P., Pless, R. & Dutcher, S. Efficient spatiotemporal analysis of the flagellar waveform of Chlamydomonas reinhardtii. Cytoskeleton 67, 56–69 (2010).

[30] Chakrabarti, B. & Saintillan, D. Spontaneous oscillations, beating patterns, and hydrodynamics of active microfilaments. Physical Review Fluids 4, 043102 (2019).

[31] Berg, H. C. & Brown, D. A. Chemotaxis in Escherichia coli analysed by three-dimensional tracking. Nature 239, 500–504 (1972).

[32] Polin, M., Tuval, I., Drescher, K., Gollub, J. P. & Goldstein, R. E. Chlamydomonas swims with two “gears” in a eukaryotic version of run-and-tumble locomotion. Science 325, 487–490 (2009).

[33] Cohen, A. E., Hastewell, A. D., Pradhan, S., Flavell, S. W. & Dunkel, J. Schrödinger dynamics and berry phase of undulatory locomotion. Physical Review Letters 130, 258402 (2023).

[34] Trefethen, L. N. Approximation theory and approximation practice, extended edition (SIAM, 2019).

[35] Arts, L. P. & van den Broek, E. L. The fast continuous wavelet transformation (fCWT) for real-time, high-quality, noise-resistant time–frequency analysis. Nature Computational Science 2, 47–58 (2022).

[36] Yoshimura, K., Shingyoji, C. & Takahashi, K. Conversion of beating mode in Chlamydomonas flagella induced by electric stimulation. Cell motility and the cytoskeleton 36, 236–245 (1997).

[37] Bottier, M., Thomas, K. A., Dutcher, S. K. & Bayly, P. V. How does cilium length affect beating? Biophysical Journal 116, 1292–1304 (2019).

[38] Son, K., Brumley, D. R. & Stocker, R. Live from under the lens: exploring microbial motility with dynamic imaging and microfluidics. Nature Reviews Microbiology 13, 761–775 (2015).

[39] Shapere, A. & Wilczek, F. Geometry of self-propulsion at low Reynolds number. Journal of Fluid Mechanics 198, 557–585 (1989).

[40] Hatton, R. L., Ding, Y., Choset, H. & Goldman, D. I. Geometric visualization of self-propulsion in a complex medium. Physical Review Letters 110, 078101 (2013).

[41] Pierce, C. J. et al. Dispersion relations for active undulators in overdamped environments. arXiv (2024).

[42] Cammann, J., Laeverenz-Schlogelhofer, H., Wan, K. Y. & Mazza, M. G. Form and function in biological filaments: A physicist’s review. arXiv (2025).

[43] Shiba, K. & Inaba, K. Inverse relationship of Ca^2+^-dependent flagellar response between animal sperm and Prasinophyte algae. Journal of Plant Research 130, 465–473 (2017).

[44] Lindemann, C. B. & Lesich, K. A. The mechanics of cilia and flagella: What we know and what we need to know. Cytoskeleton 81, 648–668 (2024).

[45] Cass, J. F. & Bloomfield-Gadêlha, H. The reaction-diffusion basis of animated patterns in eukaryotic flagella. Nature Communications 14, 5638 (2023).

[46] Man, Y., Ling, F. & Kanso, E. Cilia oscillations. Philosophical Transactions of the Royal Society B 375, 20190157 (2020).

[47] Camalet, S., Jülicher, F. & Prost, J. Self-organized beating and swimming of internally driven filaments. Physical Review Letters 82, 1590–1593 (1999).

[48] Wan, K. Y. Active oscillations in microscale navigation. Animal Cognition 26, 1837–1850 (2023).

[49] Camalet, S., Duke, T., Jülicher, F. & Prost, J. Auditory sensitivity provided by self-tuned critical oscillations of hair cells. Proceedings of the National Academy of Sciences 97, 3183–3188 (2000).

[50] Miano, F., Asadzadeh, S. S., Ryderheim, F., Andersen, A. & Kiørboe, T. High-speed escape jumps in haptophytes: Mechanism and triggering fluid signal. Limnology and Oceanography 69, 2846–2858 (2024).

[51] Gilbert, J. Jumping behavior in the oligotrich ciliates Strobilidium velox and Halteria grandinella, and its significance as a defense against rotifer predators. Microbial Ecology 27, 189–200 (1994).

[52] Jakobsen, H. H. Escape response of planktonic protists to fluid mechanical signals. Marine Ecology Progress Series 214, 67–78 (2001).

[53] Gu, H. et al. Magnetic cilia carpets with programmable metachronal waves. Nature Communications 11, 2637 (2020).

[54] Peerlinck, S. et al. Artificial cilia–bridging the gap with nature. Advanced Functional Materials 33, 2300856 (2023).

[55] Ribeiro, F. et al. Short depuration of oysters intended for human consumption is effective at reducing exposure to nanoplastics. Environmental Science & Technology 56, 16716–16725 (2022).

[56] Borg, I. & Groenen, P. J. Modern multidimensional scaling: Theory and applications (Springer, New York, USA, 2005).

