## Supplementary material for "Embodied behavioural complexity in a ciliated microorganism": SI text file

#### A Morphology Characterisation

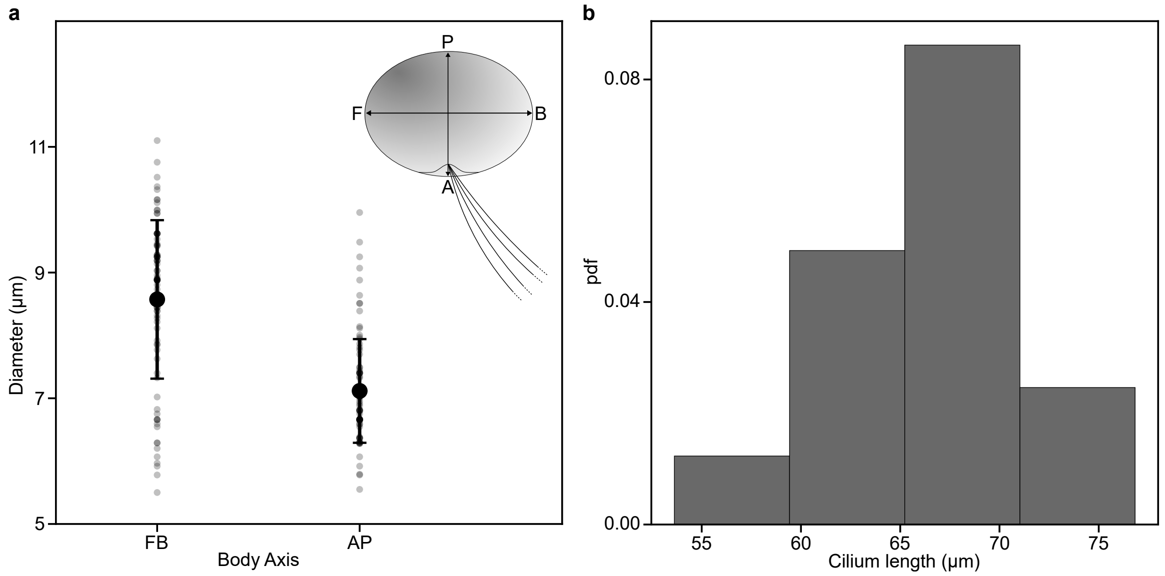


Figure 1: **a**, Diameter measurements of cell body taken from differential interference contrast microscopy images of swimming cells where the ciliary attachment point can be observed ($n_{\text{cells}} = 89$). The front-back (F-B) and anterior-posterior (A-P) axes are indicated in the cell schematic. Individual measurements are shown in grey with the mean and standard deviation in black. **b**, Histogram of cilium length measurements obtained from cilium traces of cells in their Stop state where the full length of the cilium can be observed ($n_{\text{cells}} = 14,\ n_{\text{cilia}} = 16$).

The cell body size is obtained from manual measurements in ImageJ of 89 swimming cells where the front-back (F-B) & anterior-posterior (A-P) axes of the cell can be identified based on the swimming direction and cilia attachment point (Supplementary Fig. 1a). *Pterosperma* has a typical body size of $9 \pm 1\,\mu$m (F-B) & $7.1 \pm 0.8\,\mu$m (A-P) (mean $\pm$ s.d.). The body axis definition is that of Inouye *et al.* [1].

The cilium lengths are taken from the mean traced length for 16 cilia in the Stop state. The Stop state is chosen because the entire cilium length can typically be tracked whereas while swimming the attachment point to the cell is obscured. The typical cilium length is $67 \pm 6\,\mu$m (mean $\pm$ s.d.) (Supplemental Fig. 1b).

### B Population Analysis

#### B.1 Population Trajectory Analysis & State Identification

Cells were acclimatised in image chambers for 15 min prior to population video capture. For each experiment data was captured for 2.5 min at 100 fps under brightfield illumination with a 10× (HC PL/0.40) dry objective.

Prior to tracking each image was resized to reduce its dimensions by $\frac{1}{2}$ and a background subtraction conducted by removing first a maximum intensity projection through the entire stack, removing any objects that do not move through the entire video, followed by a $15\,\mu$m median filtered image for each frame using ImageJ. Cell tracks were obtained using the TrackMate plugin for ImageJ [2] with a Laplacian of Gaussian detector and Kalman tracker.

Trajectory analysis and state identification was conducted using Python. Raw trajectories consisted of 2D cartesian coordinates $\mathrm{r}(t_n)$ at time intervals $\Delta t = t_{n+1} - t_n$ for frames $n = 1, 2, ...N$.

Swimming velocities were computed using a second order savitzky-golay filter with a 0.19 s centred window and a first derivative. This implementation first approximates each window of the trajectory with the chosen order polynomial and then uses this polynomial to compute the derivative at the given point. Cells residing in their Stop state were identified as those with instantaneous speeds $v(t) \leq 20\,\mu\mathrm{m\ s}^{-1}$ in accordance with single cell studies. Within periods of the Stop behaviour a cell could be disturbed by another passing cell without causing it to transition to the Swim state. These short periods $t \lesssim 0.5$ s where the cell speed jumps before returning to Stop state speeds were re-identified as Stop states and are the source of the small number of higher swimming speeds observed in Fig. 1g.

From the subset of the trajectory categorised as the cell swimming, periods of reorientation were identified by the change in the swimming direction. We define the turning angle $\Delta\phi = \phi(t + \Delta t) - \phi(t)$ as the difference between the swimming angle $\phi$ between the instantaneous velocity vector and the lab frame horizontal. Reorientations are then identified by having the cosine of the turning angle $\cos\Delta\phi(t) = \hat{v}(t) \cdot \hat{v}(t + \Delta t) < 0.99$ as well as a minimum in the velocity as observed in single cell trajectories of reorientations (Supplementary Fig. 2).

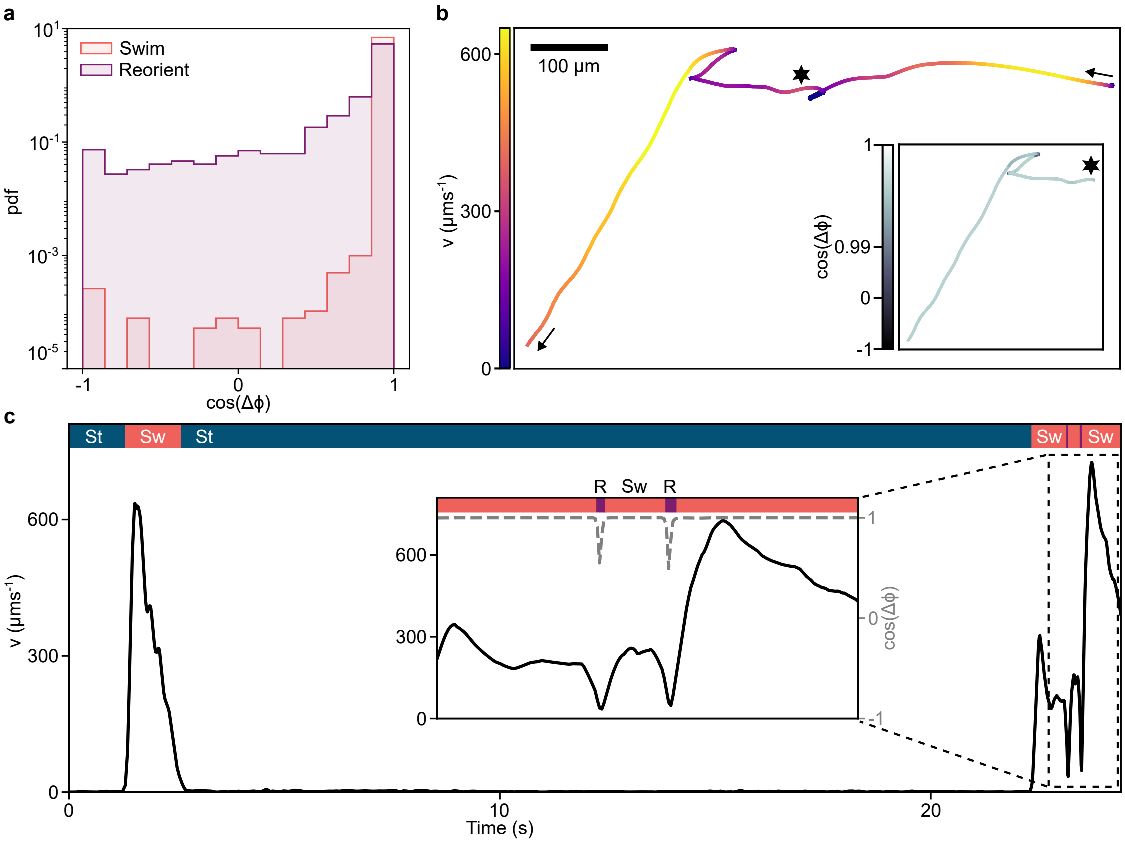


Figure 2: **a**, Histogram of the cosine of the turning angle $\cos \Delta\phi$ in the Swim and Reorient states. **b**, Example cell trajectory performing each of the three behavioural macrostates. Main panel: colour indicates the cell body speed $v$ black arrows indicate direction of travel, black star represents start of inset. Inset panel: colour indicates $\cos \Delta\phi$. **c** Cell body speed $v$ and cosine angle $\cos \Delta\phi$ profiles corresponding to the trajectory in **b**. Black line indicates $v$, dashed grey line in inset indicates $\cos \Delta\phi$, colour bar indicates the cell swimming state: Swim (coral), Reorient (purple), Stop (teal).

#### B.2 State Transition Statistics

To extract statistics that characterise our three behavioural macrostates and transitions between them the underlying process is modelled as a continuous-time Markov chain. As such we expect each state to be 'memoryless', the probability of the cell being in state $i$ at time $t$ depends only on the state it was in at $t - \Delta t$.

The Markov model used here is implemented using the R msm package [3]. The state of a cell at time $t$ denoted $S(t)$ will evolve according to the transition rates:

$$q_{jk}(t) = \lim_{\delta t \to 0} \frac{P(S(t + \Delta t) = k | S(t) = j)}{\Delta t} \tag{1}$$

The transition rate matrix $Q(t)$ is defined such that $q_{jj} = -\Sigma_{k \neq j} q_{jk}$. For the Markov property to hold the state lifetimes should be exponentially distributed as $e^{-\frac{1}{q_{jj}} t}$, giving the mean lifetime of state $S_j$ as $< \tau_j >= -1/q_{jj}$. In our implementation the a priori assumption is that $Q$ allows all transitions with equal rates with the exception of Stop $\leftrightarrow$ Reorient transitions where none were observed and are prescribed a rate of 0.

The state probability matrix $P(t) = e^{Qt}$ contains the elements $p_{jk}$ that represent the probability of being in state $k$ at time $t$ given the cell was in state $j$ at time $t - \Delta t$. By minimising the negative log-likelihood calculated from $P(t)$ the transition intensities are estimated.

For a Markov chain containing a finite number of recurrent states with no periodicity in the way transitions occur between these states a unique stationary probability distribution $P(t \to \infty) = \pi$ should emerge at long times. $\pi = (P_{Stop}, P_{Swim}, P_{Reorient})$ identifies the frequency with which we would expect to find the system in each of the three states.

The Markov model was applied to state dynamics extracted from 2096 cell trajectories including 3947 transitions across three experiments. Trajectory lengths ranged from $1 - 150\,\mathrm{s}$.

Application of this Markov framework yields final transition rates (mean ± s.e.):

$$Q = \begin{array}{c} \\ \text{Stop} \\ \text{Swim} \\ \text{Reorient} \end{array} \begin{array}{c} \begin{array}{ccc} \text{Stop} & \text{Swim} & \text{Reorient} \end{array} \\ \begin{pmatrix} -0.0174 \pm 0.0005 & 0.0174 \pm 0.0005 & 0 \\ 0.50 \pm 0.01 & -0.71 \pm 0.02 & 0.21 \pm 0.01 \\ 0 & 24 \pm 1 & -24 \pm 1 \end{pmatrix} \end{array}$$

And associated mean state lifetimes:

$$(\langle \tau_{\text{Stop}} \rangle, \langle \tau_{\text{Swim}} \rangle, \langle \tau_{\text{Reorient}} \rangle) = (58 \pm 2, 1.42 \pm 0.03, 0.041 \pm 0.002) \tag{2}$$

The state probabilities converge to the steady state values as $t \to \infty$ giving:

$$(P_{\text{Stop}}, P_{\text{Swim}}, P_{\text{Reorient}}) = (0.966 \pm 0.002, 0.033 \pm 0.001, 0.00030 \pm 0.00005) \tag{3}$$

As a more intuitive interpretation of the transition dynamics we can also extract the probability that the next state is $S_k$ given the initial state was $S_j$ as $\rho_{jk} = -q_{jk}/q_{jj}$ yielding $(\rho_{\text{St}\to\text{Sw}}, \rho_{\text{Sw}\to\text{St}}, \rho_{\text{Sw}\to\text{R}}, \rho_{\text{R}\to\text{Sw}}) = (1, 0.70 \pm 0.04, 0.30 \pm 0.04, 1)$ noting that $\rho_{\text{Sw}\to\text{St}} + \rho_{\text{Sw}\to\text{R}} = 1$

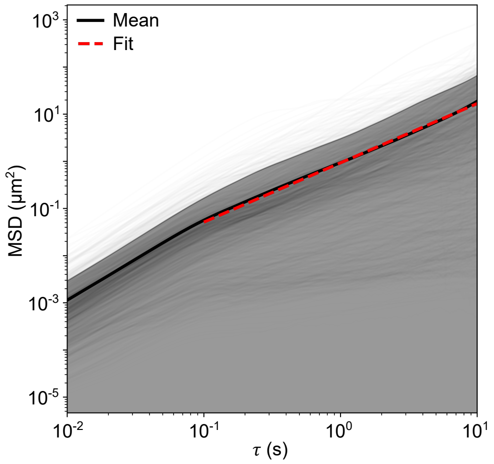


Figure 3: Mean squared displacement (MSD) vs lag time $\tau$ in the Stop state. Grey lines indicate the 1314 trajectories selected with lengths longer than 10 s. The mean MSD is illustrated in black with shading over one standard deviation. A fit (red dashed line) is performed over the range $\tau = 0.1 - 10$ s.

##### B.3 Mean squared displacement in the Stop state

From the identified population trajectories we separate the Stop data as per the initial thresholding outlined in Section B.1. We compute the lag time mean squared displacement $\text{MSD} = \frac{1}{N-\tau}\sum_{t=0}^{N-\tau-1} |\mathbf{r}(t_n+\tau) - \mathbf{r}(t_n)|^2$ where there are $N$ frames in total and $\tau = 1, 2, ...N-1$ is the temporal lag (Supplementary Fig. 3). We then fit an anomalous diffusion equation to the average MSD from all cells which, in two dimensions, takes the form $\text{MSD} = 4Dt^{\alpha}$ where $D$ is the effective diffusion coefficient and $\alpha$ is the anomalous exponent. $\alpha = 1$ indicates diffusion while $\alpha > 1$ corresponds to super-diffusive behaviour.

We focus on an intermediate range of times where many trajectories are identified, selecting trajectories of at least 10 s in length. From these 1314 trajectories we compute the mean MSD and fit over the range $\tau = 0.1 - 10$ s to obtain a value of $D = 0.2\,\mu\text{m}^2\ \text{s}^{-1}$ and $\alpha = 1.3$. As such the Stop dynamics are slightly super-diffusive.

#### C Cilium tracking

Extraction of the cilium tracks is performed using MATLAB® following the protocol of Walker *et al.* [4] and illustrated in Supplementary Fig. 4a.

Videos of cells are processed by the subtraction of a mean intensity projection (excluding Stop state data) and application of a Frangi vesselness filter prior to thresholding.

Binary images are skeletonised and the cilium centreline extracted using a threshold on the medial axis transform (MAT) (Supplementary Fig. 4a). The medial axis transform is the multiplication of the skeletonised object with a distance transform computed on the binary image measuring the 'thickness' of the object.

Rather than typical spline interpolation to smooth cilium traces we project the raw traces on to Chebyshev polynomials of the first kind $T_n(s)$ [5]. The ciliary waveforms $\mathbf{r}(s,t) = (x(s,t), y(s,t))$ can be described purely by the leading order Chebyshev coefficients $\hat{\mathbf{r}}_n$ up to degree $N$. Given a set of segmented cilium centre line pixels $\{\mathbf{p}_k\}$ ordered

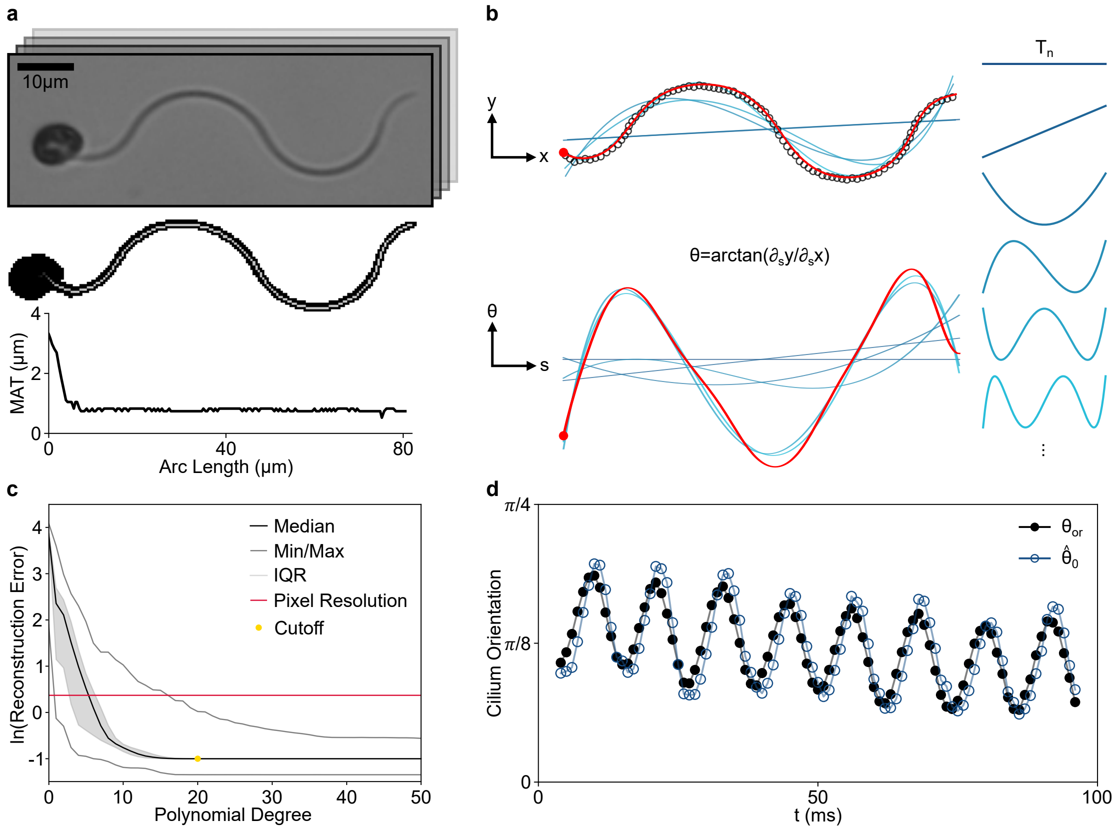


Figure 4: **a**, Extraction of cilium centreline from video frames. Top: differential interference contrast microscopy image of cell. Middle: binarised cell overlaid with medial axis transform (MAT). Bottom: MAT profile from cell body to cilium tip illustrating separation between cell body and cilium 'thickness'. **b**, Schematic of the superposition of Chebyshev modes that reconstructs the cilium shape shown for the first 6 modes. Top: Cartesian space representation. Bottom: Tangent space representation. Black circles denote the raw trace sampled every 2 pixels, blue lines denote the superposition of an increasing number of modes approaching the cilium shape, red line denotes the final smoothed cilium trace. **c**, Root mean square reconstruction error as a function of the number of Chebyshev modes used. The median error is shown in black with the interquartile range (IQR) shaded, the minimum and maximum error across across the waveform dataset is displayed in grey. Red line indicates the pixel resolution of 0.370 $\mu$m/pixel. $\mathrm{n}_{\mathrm{cutoff}} = 20$ modes are used in the final tangent angle parametrisation. **d**, Cilium orientation as a function of time. Black filled circles indicate the angle between the line passing through the start and end point of the cilium and the x-axis $\theta_{or} = \arctan\left(\frac{y[L]-y[0]}{x[L]-x[0]}\right)$ blue unfilled circles indicate the $0^{th}$ mode of the Chebyshev decomposition.

consistently with the arc length ordering, $k_1 < k_2 \implies s_1 < s_2$, we can fit the coefficients in the Chebyshev expansion as the minimisation of a least squares problem:

$$\{\hat{\mathbf{r}}_n^*\}_{n=0}^N = \min_{\hat{\mathbf{r}}_n} \sum_{k=1}^{K} \left\| \mathbf{p}_k - \sum_{n=0}^{N} \hat{\mathbf{r}}_n T_n(q_k) \right\|_2^2 \tag{4}$$

where the $q_k$ are approximated from the data using the discrete arc length approximation:

$$L_d = \sum_{k=1}^{K-1} \|\mathbf{p}_{k+1} - \mathbf{p}_k\|_2,\ q_{k+1} - q_k = \frac{2}{L_d}\|\mathbf{p}_{k+1} - \mathbf{p}_k\|_2$$

with $q_1 = -1$ and $q_N = 1$. Since we assume that the points are ordered this choice of $q_k$ amounts to a specific but not arc length parametrisation of the curve. Defining the matrix $\mathcal{T}_{nk} = T_k(t_n)$ and stacking all the coordinates of $\mathbf{p}_k$ into length $K$ vectors $\mathbf{x}$ and $\mathbf{y}$ respectively, we can write the minimisation problem in matrix form, separately for each dimension:

$$\min_{\hat{\mathbf{x}}} \|\mathcal{T}\hat{\mathbf{x}} - \mathbf{x}\|_2^2, \qquad \min_{\hat{\mathbf{y}}} \|\mathcal{T}\hat{\mathbf{y}} - \mathbf{y}\|_2^2. \tag{5}$$

where $\hat{\mathbf{r}}_n = (\hat{x}_n, \hat{y}_n)$. In general this problem will be ill-posed for data not sampled at the Chebyshev points and when $n \sim k$. However, the known spectral decay condition for the coefficients of smooth functions provides an additional constraint $\sum_{n=0}^N |e^{\rho n}\hat{x}_n|^2 = \|E\hat{\mathbf{x}}\|^2 \leq 4NM^2$ where $M$ is the maximum absolute value of a function over its complex continuation and $\rho$ is related to the smoothness of the representation [6]. Solving Equation (5) subject to this constraint corresponds to a trust-region least squares problem that can be solved using standard optimisation techniques [7]. We estimate $M$ from the data to be fit and $\rho$ is a free parameter that controls the smoothness of the function approximation. We find that the results are not sensitive to choices of $M$ provided it is chosen large enough. Additionally since we will need to calculate derivatives of the resulting expansion we add a second order derivative total variation regularisation to the problem $\int_{-1}^1 \mathrm{d}s\, (x''(s))^2$ that can be expressed in terms of the coefficients as $\|D_2 I D_2 \hat{\mathbf{x}}\|$ where $D_2$ is the second order Chebyshev differentiation matrix and I the Chebyshev integration matrix. We optimise $\hat{\mathbf{x}}$ and $\hat{\mathbf{y}}$ separately using the combined optimisation:

$$\min_{\|E\hat{\mathbf{x}}\|^2 \leq 4NM^2} \left( \|\mathcal{T}\hat{\mathbf{x}} - \mathbf{x}\|_2^2 + \lambda \|D_2 I D_2 \hat{\mathbf{x}}\| \right) \tag{6}$$

where $M$ is determined from the data, $\lambda = 0.00005$ and $\rho$ is varied to provide the smoothest possible function with a root mean squared error less than a size of a pixel on average. Due to regularisation we fit $N = 125$ coefficients. While these are not all necessary to describe the shape they are important for being able to calculate smooth, high quality derivatives which is important for re-parametrisation and computing the angular representation.

The coefficients $\hat{\mathbf{x}}$ and $\hat{\mathbf{y}}$ give a smooth representation for $\mathbf{r}(q, t)$ but $q$ is not necessarily proportional to an arc length parametrisation which requires $|\partial_q \mathbf{r}(q, t)| = 1$. We re-parametrise the curve as follows: first we calculate the length of the representation $L = \int_{-1}^1 \mathrm{d}q\, |\partial_q \mathbf{r}(q, t)|$ where the derivative is calculated using the Chebyshev representation. At a given time point the normalised arc length value at $q$ is given by:

$$s(q) = -1 + \frac{2}{L} \int_{-1}^{q} \mathrm{d}q'\, |\partial_q \mathbf{r}(q)| \tag{7}$$

Our goal is to determine what value of $q_k^c$ correspond to the arc length sampled at the Chebyshev nodes $s_k^c = \cos \pi(k - 1/2)/N = s(q_k^c)$ for $k = 0, 1, \ldots, N - 1$. This corresponds to a root finding problem on the function $S_k(q) = s(q) - s_c^k$ which can be solved efficiently using Newton's method where derivatives are computed using the Chebyshev representation. We can then sample the representation of $\mathbf{r}(q, t)$ at $q_k^c$ and apply the discrete cosine transform (DCT) to get a new set of coefficients $\tilde{\mathbf{r}}_n$ that correspond to the arc length paramterised representation of the center line in Cartesian coordinates, $\mathbf{r}(s, t) = \sum_n T_n(s)\tilde{\mathbf{r}}_n(t)$.

This representation allows a smooth transformation into the frame of reference of the cilium by computing the tangent angle:

$$\theta(s, t) = \arctan(\partial_s y(s, t)/\partial_s x(s, t)) = \sum_{n=0}^{N} T_n(s)\hat{\theta}_n(t) \tag{8}$$

The arc length and Cartesian space representations of the cilium can be recovered as:

$$\begin{bmatrix} x(s,t) \\ y(s,t) \end{bmatrix} = x(0, t) + \frac{L}{2}\int_{-1}^{s} ds' \begin{bmatrix} \cos\theta(s', t) \\ \sin\theta(s', t) \end{bmatrix} \tag{9}$$

where $L$ is the total trace length. From the definition of the coefficients:

$$\hat{\theta}_n(s) = \frac{k_n}{\pi}\int_{-1}^{1} \mathrm{d}s\, \frac{\theta(s, t)}{\sqrt{1 - s^2}} T_n(s) \tag{10}$$

where $k_0 = 2$ and $k_n = 1$ otherwise. We see that the $0^{\text{th}}$ coefficient corresponds to a weighted average orientation since $T_0(s) = 1$:

$$\hat{\theta}_0(s) = \frac{2}{\pi}\int_{-1}^{1} \mathrm{d}s\, \frac{\theta(s, t)}{\sqrt{1 - s^2}}. \tag{11}$$

Similarly the $1^{\text{st}}$ coefficient represents a weighted average of the signed curvature:

$$\hat{\theta}_1(s) = \frac{1}{\pi}\int_{-1}^{1} \mathrm{d}s\, \frac{s\theta(s, t)}{\sqrt{1 - s^2}} = \frac{1}{\pi}\int_{-1}^{1} \mathrm{d}s\, \partial_s\theta(s, t)\sqrt{1 - s^2} = \frac{1}{\pi}\int_{-1}^{1} \mathrm{d}s\, \kappa(s)\sqrt{1 - s^2} \tag{12}$$

where $\kappa(s)$ is the signed curvature.

We define an approximate wavelength for the $n$th Chebyshev polynomial by $\frac{1}{2}\lambda_n = \frac{L}{2(n-1)}\sum_{l=0}^{n-2} \cos(\frac{\pi l}{n} + \frac{\pi}{2n}) - \cos(\frac{\pi(l+1)}{n} + \frac{\pi}{2n})$ which is the mean distance between the extrema scaled by the average cilium length. The factor of 2 in the length scaling is due to the natural interval for Chebyshev polynomials being $[-1, 1]$.

### D Reorientation stereotypy

To characterise reorientations we use body trajectories from 14 cells to identify the reorientation angles and times (Supplementary Fig. 5a). The start of the reorientation can be identified by a peak in the squared turning angle, $(\Delta\phi)^2$ (Supplementary Fig. 5b). As the cell emerges from the reorientation it relaxes back into the Swim state which can

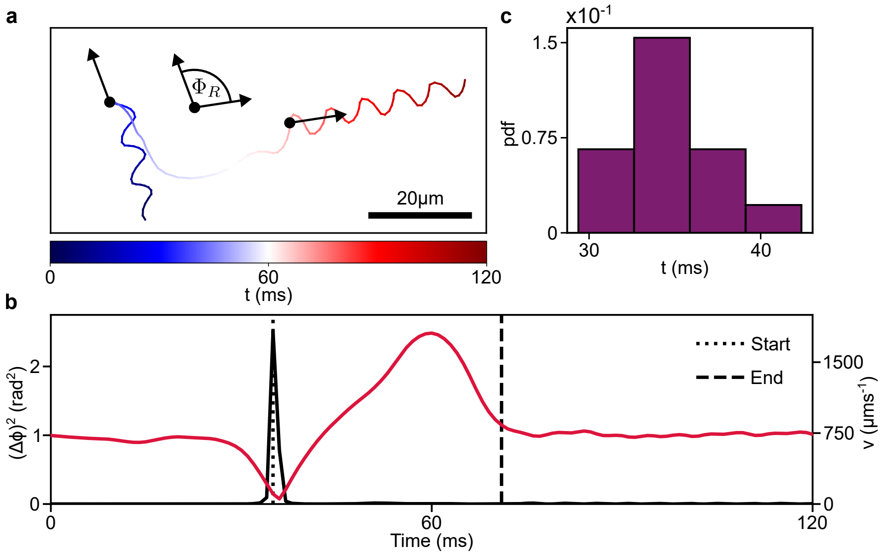


Figure 5: **a**, Cell body trajectory during a reorientation. Colour indicates time, black arrows indicate the incoming and outgoing direction of the cell at the start and end of the reorientation. **b**, Corresponding squared turning angle $(\Delta\phi)^2$, and cell body speed $v$. Black dotted line indicates the start of the reorientation, black dashed line indicates the end of the reorientation. **c**, Histogram of reorientation times calculated for $n = 14$ reorientations, yielding a reorientation time $\tau_R = 36 \pm 3\,\mathrm{ms}$ (mean ± s.d.)

be observed in the decay of the body speed into steady swimming (Supplementary Fig. 5b). We identify the end point of the reorientation where this deceleration drops to zero within a tolerance of $\frac{1}{2}\frac{dv}{dt}\big|_{\max}$. We can then calculate the reorientation time $\tau_R$.

Having identified start and end points of the reorientation we identify the incoming and outgoing swimming directions and compute the angle between these as the reorientation angle $\Phi_R$ (Supplementary Fig. 5a). We apply a backwards (forwards) facing window of 10 ms to compute the mean incoming (outgoing) swimming direction.

From our reorientation dataset it was possible to accurately identify the start and end points of the reorientation while also tracking the cilia throughout the reorientation for 4 cells. We identify each waveform trace with a phase $\psi(t) = \frac{2\pi}{\tau_R}t$ and discretise these into 8 bins, where $t \in [0, \tau_R]$. The bin average tangent waveform is then computed, the cartesian space representation of this is illustrated in Figure 2h.

### E Wavelet analysis & dispersion relation

To generate the wavelet vectors illustrated in Fig. 2f & g a continuous wavelet transform was performed on each of the Chebyshev mode amplitudes independently up to $N = 20$. Wavelet analysis was performed via the Fourier transform using a Morlet wavelet with characteristic frequency of $2\pi$ [8, 9]. The initial and final 20 frames are cropped from the resulting transform to remove boundary artifacts. We stack the wavelets corresponding to Chebyshev modes of increasing wavenumber to generate the wavelet vector representation.

We apply the analysis described to a large high-resolution dataset of single-cell ciliary dynamics consisting of 219,368 frames of ciliary bundle shapes from a total of 125 cells incorporating all transitions observed in the population level transition network (Fig. 1i). In the Stop state this relies on the fact that before the cilia fully unbundle the cell has settled into the very low frequency beating characteristic of this state and in many cases the cilia never fully unbundle allowing us to capture transitions to swimming.

To compute the $f$-$k$ dispersion we average over the set of wavelet vectors for our entire dataset. This generates a single column in our wavelet vector representation capturing, for each Chebyshev mode, a mean frequency profile. We construct the $f$-$k$ space by stacking these frequency profiles at increasing wavenumber. This is plotted with a linear frequency scale in Supplementary Fig. 6a illustrating the approximately linear dispersion relation, $f \sim k$. This scale suppresses the location of the low frequency Stop state dynamics hence the choice of a logarithmic scale in Fig. 3h.

An alternative representation (Supplementary Fig. 6b) shows clearly the presence of the 'steps' in the dispersion plot. Here we see the association of peaks in the frequency spectrum with different sets of wavenumbers. By selecting the largest amplitude peak in each case we identify the typical frequency observed in each band of the diagonal branch of the dispersion plot. This yields frequency bands at approximately 37, 88, 184, and 265 Hz.

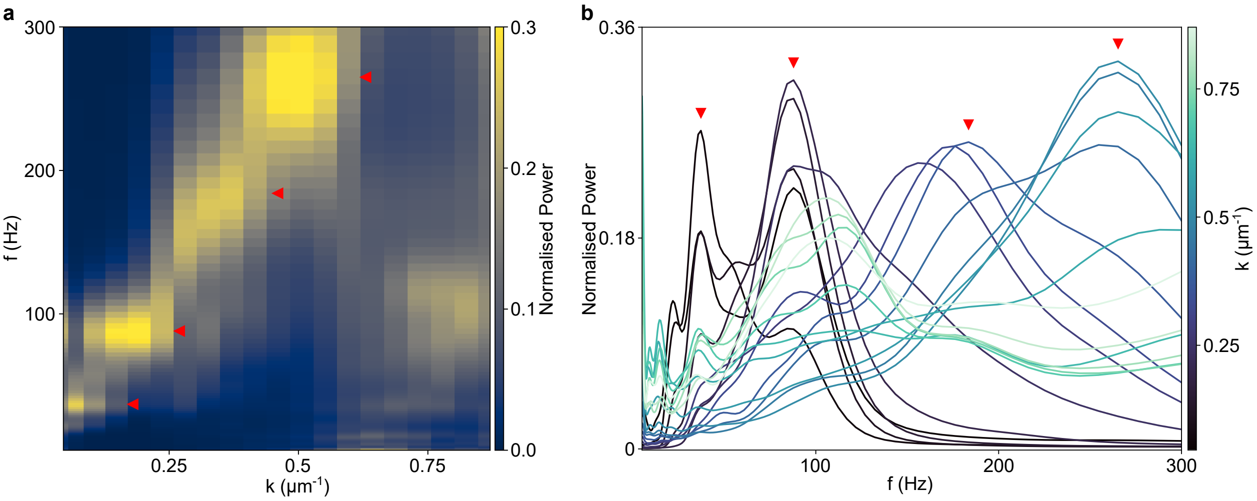


Figure 6: **a**, Dispersion relation with linear frequency scale. Frequency peaks in **b** are indicated by red arrows. **b**, Frequency spectrum at each wavenumber. Four peaks are observed at varying $k$ indicated by red arrows at $f = 37,\ 88,\ 184,\ 265\,\mathrm{Hz}$

### F Multidimensional scaling embedding

Multidimensional scaling (MDS) is performed on a distance matrix computed from the Euclidean distances between instantaneous wavelet vectors. The distances are calculated between the normalised wavelet amplitudes $d_{ij} = \left|\hat{\mathbf{W}}(t_i) - \hat{\mathbf{W}}(t_j)|\right|$.

When performing the MDS the wavelet data is randomly segmented into three bins with 1666 time points shared between the bins. MDS is then performed separately on each bin. As MDS is only defined up to an affine transformation, the resulting MDS coordinates are then aligned by fitting an affine transform between the overlapping points in each bin to produce a combined MDS embedding [10, 11].

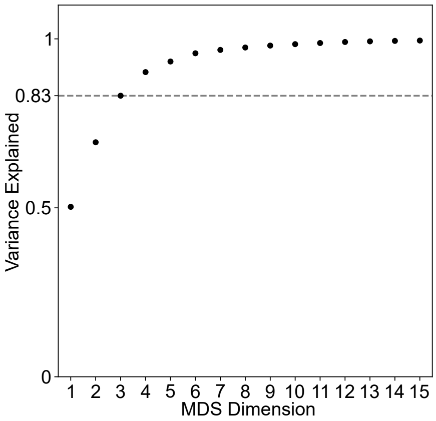


Figure 7: Variance explained by the MDS dimensions.

The variance explained by the MDS dimensions (Supplementary Fig. 7) is computed as the normalised sum of the squared eigenvalues of the distance matrix. In Figure 4b we illustrate this low dimensional embedding in three dimensions which captures 83% of the variance in the wavelet vector data. 6 dimensions are required to capture $> 95\%$ of the variance.

In Supplementary Fig. 8 we colour a reduced manifold by three key physical parameters: the cell speed, cilium beat frequency & wavenumber, as well as MDS 4. For the frequency and wavenumber we can construct an instantaneous $f$-$k$ space as described for the dispersion plots but now taking a single time point and hence column of the wavelet vector. To assign a single value to this we take the mean value in the instantaneous $f$-$k$ space.

We exclude the Stop state data in these representations to focus on the diverse dynamics during swimming. In addition, as observed in the dispersion plot (Figure 4a) the Stop dynamics combine a low $k$ component that corresponds to an effectively static underlying curve in the cilium waveform and the high $k$, low $f$ ciliary oscillations. Thus taking the instantaneous mean frequency and wavenumber for the Stop state would capture values that sit between these two features of the waveform dynamics. In Supplementary Video 3, we use an alternative criteria to select the $f$, $k$ values for the Stop-Swim trajectory. We use an exponential 'soft-max' weighting $e^{\sigma x}/\sum_x e^{\sigma x}$ with $\sigma = 25$ applied to the $f$-$k$ space wavelet amplitudes before averaging. This allows us to select the dynamic component of the Stop state waveform and thus follow its trajectory on the dispersion plot during these transitions.

In Supplementary Fig. 8, we observe various trends in the physical parameters. We find the cell reduces its swimming speed $v$ as it transitions towards the Stop state. This process involves the $f$ and $k$ ramping discussed and observable in the co-location of reduced swimming speeds with increasing frequency and wavenumber. In both frequency and wavenumber we see the growth of each parameter as we trace a path from the peak at positive MDS-1 & negative MDS-2 along the manifold envelope. This growth is more discrete in $f$ mirroring the steps observed in the dispersion plot.

We also map the value of MDS 4 in Supplementary Fig. 8d. The clear structure

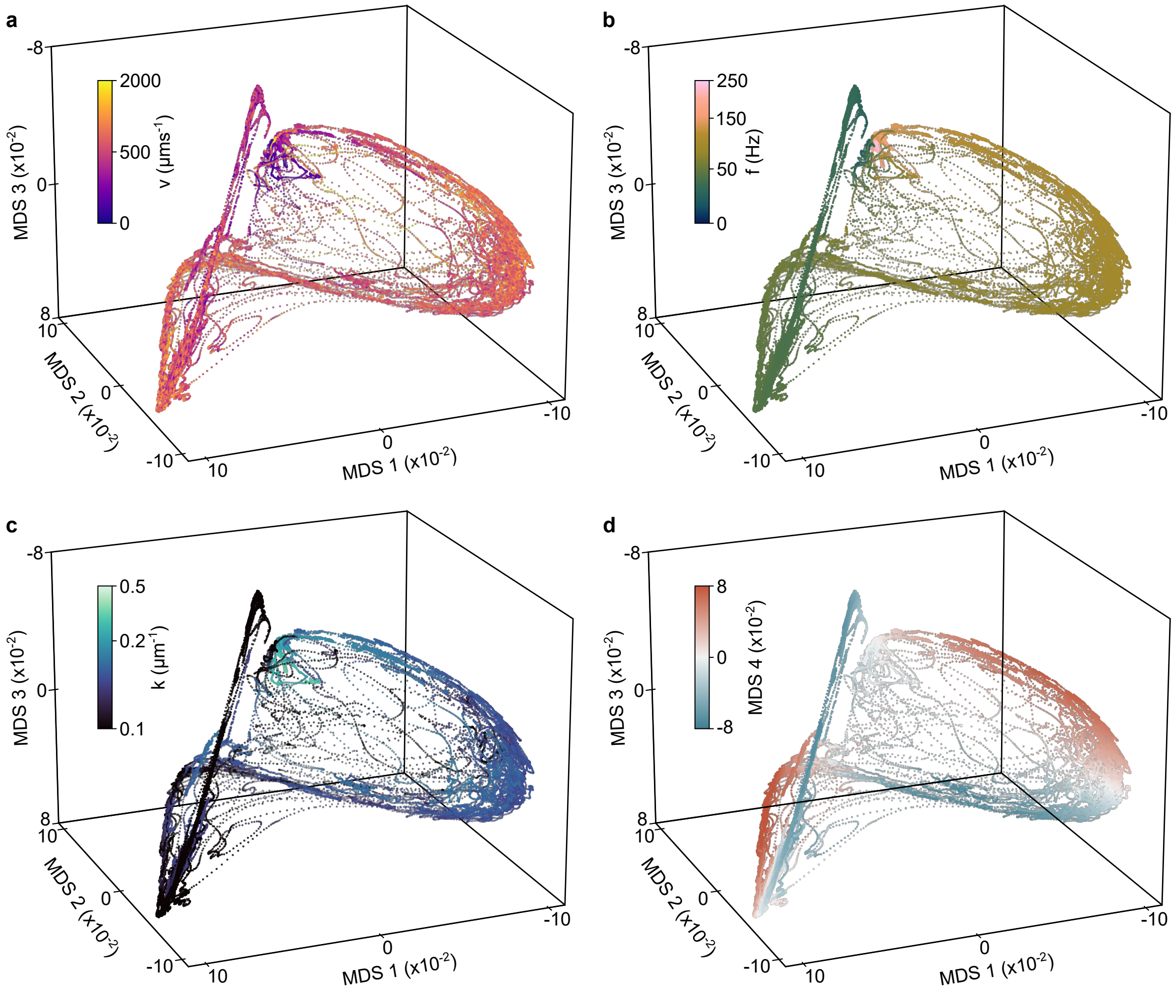


Figure 8: Behavioural manifold, excluding the Stop data, coloured by instantaneous cell speed (**a**), mean cilium frequency (**b**), mean cilium wavenumber (**c**) and MDS 4 (**d**).

in MDS 4 illustrates the noted higher dimensionality of the manifold than the three dimensions illustrated.

### References


[1] Inouye, I., Hori, T. & Chihara, M. Absolute configuration analysis of the flagellar apparatus of *Pterosperma cristatum* (Prasinophyceae) and consideration of its phylogenetic position. *Journal of Phycology* **26**, 329–344 (1990).

[2] Tinevez, J.-Y. *et al.* Trackmate: An open and extensible platform for single-particle tracking. *Methods* **115**, 80–90 (2017).

[3] Jackson, C. Multi-state models for panel data: The msm package for R. *Journal of Statistical Software* **38**, 1–28 (2011).

[4] Walker, B. J., Ishimoto, K. & Wheeler, R. J. Automated identification of flagella from videomicroscopy via the medial axis transform. *Scientific Reports* **9**, 5015 (2019).

[5] Mason, J. C. & Handscomb, D. C. *Chebyshev polynomials* (Chapman and Hall/CRC, 2002).

[6] Trefethen, L. N. *Approximation theory and approximation practice, extended edition* (SIAM, 2019).

[7] Nocedal, J. & Wright, S. J. *Numerical optimization* (Springer, 2006).

[8] Arts, L. P. & van den Broek, E. L. The fast continuous wavelet transformation (fCWT) for real-time, high-quality, noise-resistant time–frequency analysis. *Nature Computational Science* **2**, 47–58 (2022).

[9] Addison, P. S. Introduction to redundancy rules: the continuous wavelet transform comes of age. *Philosophical Transactions of the Royal Society A* **376**, 20170258 (2018).

[10] Borg, I. & Groenen, P. J. *Modern multidimensional scaling: Theory and applications* (Springer, New York, USA, 2005).

[11] Delicado, P. & Pachón-García, C. Multidimensional scaling for big data. *Advances in Data Analysis and Classification* 1–22 (2024).
